# Bone marrow microenvironment signatures associate with patient survival after guadecitabine and atezolizumab therapy in HMA-resistant MDS

**DOI:** 10.1101/2024.11.08.622670

**Authors:** H. Josh Jang, Guillermo Urrutia, Andreas Due Orskov, Hyeon Jin Kim, Seth A. Nelson, Anh Van Nguyen, Hyein Lee, Ryan S. Burgos, Benjamin K. Johnson, Marc Wegener, Katelyn Becker, Marie Adams, Rachael Sheridan, Zachary H. Ramjan, Scott A. Givan, Caitlin C. Zebley, Benjamin A. Youngblood, Jean-Pierre J. Issa, Michael J. Topper, Stephen B. Baylin, Maria R. Baer, Timothy J. Triche, Casey L. O’Connell, Kirsten Gronbaek, Peter A. Jones

## Abstract

Almost 50% of patients with myelodysplastic syndrome (MDS) are refractory to first-line hypomethylating agents (HMAs), which presents a significant clinical challenge considering the lack of options for salvage. Past work revealed that immune checkpoint molecules on peripheral myeloblasts and immune cells are up-regulated after HMA treatment. Therefore, we conducted a Phase I/II clinical trial combining guadecitabine (an HMA) and atezolizumab (an immune checkpoint inhibitor) to treat HMA-relapsed or refractory (HMA-R/R) MDS patients. This combination therapy showed median overall survival of 15.1 months relative to historical controls (4-6 months). Here, we profiled the cell composition and gene expression signatures of cells from bone marrow aspirates from trial participants with short-term (<15 months) or long-term (>15 months) survival at single-cell resolution. Long-term survivors showed a significant reduction of immunosuppressive monocytes, and an expansion of effector lymphocytes after combination therapy. Further immune profiling suggests that gamma delta T cell activation through primed dendritic cells was associated with global interferon activation in the bone marrow microenvironment of long-term survivors. Short-term survivors exhibited elevated inflammation and senescence-like gene signatures that were not resolved by combination therapy. We propose that distinct bone marrow microenvironment features, such as senescence-associated inflammation or immunosuppressive monocyte presence, could improve patient stratification for HMA and immunotherapy combinations in HMA-R/R MDS patients.

## Introduction

Myelodysplastic syndromes (MDS) are characterized by the aberrant or disease-initiating hematopoietic stem and progenitor cells (HSPCs) in the bone marrow, which contribute to failed hematopoiesis and progression to acute myeloid leukemias^1–4^. Hypomethylating agents (HMAs) are the current standard of care for MDS patients, but ∼50% of MDS patients do not respond or progress after response to HMA therapy^5,6^. These HMA-relapsed or -refractory (HMA-R/R) MDS patients only have a median survival of ∼4-6 months^7–10^, so there is a significant clinical need to design evidence-based follow-up treatments. HMA treatment is known to epigenetically reactivate transposable elements in cancer cells, where the resulting double-strand RNAs trigger the “viral mimicry” response characterized by the up-regulation of interferon responses pathways^11–13^. The viral mimicry activation in the CD34+ bone marrow cells (which comprise various healthy and malignant HSPCs) of treatment-naïve MDS patients has been previously associated with clinical response after HMA therapy^13^. However, we and others reported that HMA therapy could also reduce tumor immunogenicity by up-regulating immunosuppressive checkpoint proteins, such as programmed cell death protein (PD-1) and programmed death-ligand 1 (PD-L1), in CD8 T cells^14^ and malignant CD34+ myeloblasts^15^, respectively. We therefore hypothesized that adding immune checkpoint inhibitors (ICIs) to HMA therapy could reverse therapy resistance through the restimulation of effector immune cells and consequential malignant myeloid cell elimination^16–18^. We recently concluded a Phase I/II study combining a HMA (guadecitabine) with PD-L1 inhibitor (atezolizumab) in HMA-R/R MDS patients^19^ and found that the combination therapy extended patient median overall survival to 15.1 months, with a subset of patients achieving a median survival of 48.9 months. This prompted us to investigate the biological mechanisms associated with long-term patient survival.

Past correlative studies have primarily focused on characterizing genetic and molecular changes in the malignant CD34+ myeloblasts to identify tumor-cell-intrinsic mechanisms that associate with clinical response^5,7,20–23^. Clinical associations were often weak or absent, suggesting that tumor-extrinsic mechanisms or dynamics of the bone marrow microenvironment, might be contributing factors to clinical efficacy. The bone marrow microenvironment of MDS patients reflects the intricate interactions between healthy or malignant CD34+ cells and stromal as well as immune cells, which collectively modulate hematopoiesis. Single-cell profiling technologies now enable us to characterize systems-level bone marrow cellular composition, and dissect how cancer heterogeneity and the presence of certain immune cell populations in the bone marrow microenvironment contribute to therapy success or failure. Here, we performed molecular profiling of cells from HMA-R/R patient bone marrow aspirates to determine how the tumor cell-intrinsic and -extrinsic factors, such as gene signatures of bone marrow CD34+ cells or immune cells in the microenvironment, might associate with improved survival outcomes after ICI + HMA therapy. We leveraged bulk- and single-cell sequencing technologies to identify pre-existing or treatment-induced transcriptional signatures and cellular composition differences in the bone marrow microenvironment. We observed that the bone marrow microenvironment in short-term survivors (<15 months survival) at baseline displayed elevated inflammatory response and contained cells with senescence-like signatures that were not resolved by combination therapy. In long-term survivors (>15 months survival), the bone marrow was enriched with primed dendritic cells, and *S100A8/9*-expressing myeloid precursor cells and monocytes. Combination therapy in long-term survivors reduced bone marrow HSPC and monocyte fractions, which was linked to effector lymphocyte expansion and systemic induction of interferon responses. We therefore suggest that pre-screening the bone marrow microenvironment of HMA-R/R MDS patients might allow us to strategically select those patients who will best benefit from combined HMA and ICI therapy.

## Results

### Bone marrow CD34+ cells have a viral mimicry signature that is decoupled from transposable element activation

We collected bone marrow biopsies and peripheral blood mononuclear cells (PBMCs) from short-term survivors (STS) or long-term survivors (LTS) before combination therapy, and at regular intervals thereafter (**Fig. S1A, Table S1**). First, we explored the potential link between viral mimicry activation and combination therapy efficacy. We isolated CD34+ cells from bone marrow aspirates using magnetic beads and generated bulk total RNA-seq libraries (**Methods**). As previously reported^13^, genes associated with the viral mimicry pathway were elevated in patient-derived CD34+ cells at baseline relative to those from healthy donors (**Fig. 1A**). Viral mimicry activation coincided with higher transposable element expression, specifically from intergenic Long Interspersed Nuclear Elements (LINEs) or Long Terminal Repeats (LTRs), in patient-derived CD34+ cells relative to healthy donor cells (**Fig. S1B**). After combination therapy, however, viral mimicry activation in CD34+ cells was decoupled from TE transcriptional induction in STS and LTS patients (**Fig. 1B & S1C**), which suggests that extrinsic cell-to-cell communication in the bone marrow microenvironment is likely influencing the immunogenicity of bone marrow CD34+ cells in HMA-R/R MDS patients.

**Figure 1.**
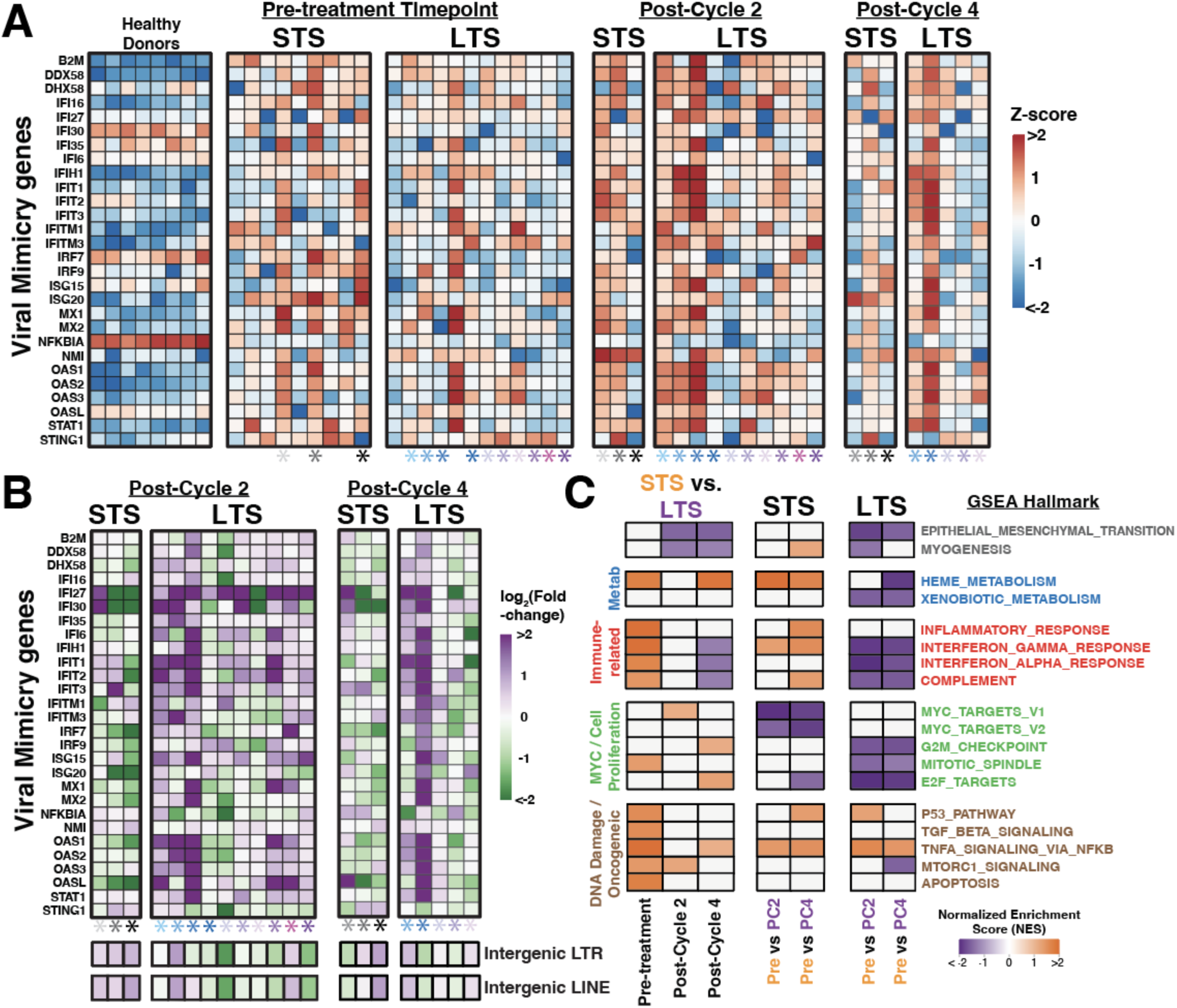
Bulk transcriptome comparisons of bone marrow CD34+ cells of HMA-R/R MDS patients. **A**-**B.** Heatmaps representing viral mimicry gene expression (**A**) or fold-change (**B**) after combination therapy in healthy donor CD34+ or patient-derived bone marrow CD34+ cells. Expression fold-changes of intergenic transposable element classes induced after treatment are illustrated below the viral mimicry fold-change heatmap. Colored asterisks represent patient ID. **C.** Hallmark Gene Set Enrichment Analysis (GSEA) comparing STS- or LTS-CD34+ cells across treatment timepoints (adjusted p-value < 0.01). Short-term survivors, STS; Long-term survivors, LTS; Pre, Pre-treatment; Post-cycle 2, PC2; Post-cycle 4, PC4; Long terminal repeat, LTR; Long interspersed nuclear element, LINE.

### Combination therapy shapes distinct bone marrow cellular dynamics in STS and LTS

To identify other tumor-intrinsic molecular pathways that might contribute to the differential survival of STS and LTS patients, we compared bone marrow CD34+ transcriptomes for immune checkpoint expressions before and after combination therapy (**Fig. S2A**). While we did not detect any statistically significant differences in immune checkpoint expression in CD34+ cells at the baseline, immunosuppressive *BLTA* and *TIGIT* expressions^24,25^ were elevated in STS-CD34+ cells, and immunostimulatory *OX40L*^24^ showed increased expression in LTS-CD34+ cells (**Fig. S2B,C**). Importantly, we did not detect differences in *CD274* (PD-L1) expression in STS- and LTS-CD34+ cells, implying that the therapeutic benefit of atezolizumab could result from modulating another PD-L1-expressing cell type in the bone marrow microenvironment.

We next queried differential gene pathway activation in these CD34+ cells using Gene Set Enrichment Analysis (GSEA)^26^. This revealed that STS-CD34+ cells at baseline showed elevated tumor necrosis factor alpha (TNFa) signaling via the nuclear factor kappa B (NFkB) pathway and inflammatory response signatures (**Fig. 1C**). Combination therapy reduced interferon gamma response and induced the MYC pathway in these cells, which likely contributed to the cancer progression in STS patients. In contrast, combination therapy increased LTS-CD34+ cell proliferation and robustly stimulated interferon response pathways to boost cancer cells’ immunogenicity in the LTS bone marrow. Relative to healthy donor CD34+ cells, patient-derived CD34+ cells also had low expression of genes from epithelial-mesenchymal transition (EMT) and collagen-containing extracellular matrix (ECM) pathways at baseline, which were restored after combination therapy specifically in LTS-CD34+ cells (**Fig. S3A-C**). While CD34 is a common HSPC marker, CD34 is also expressed in bone marrow mesenchymal stromal cells and endothelial progenitor cells^27^. Thus, these bulk transcriptomes likely represent an admixture of various CD34+ cell types.

Mesenchymal stromal cells are the primary producers of collagen in bone marrow^28,29^, and the increase in EMT and collagen-related gene expression in CD34+ cells after therapy could reflect therapy-induced expansion or recovery of MSCs in the bone marrow^30^. Indeed, in the LTS bone marrow after combination therapy, the HSPC-specific gene expression was reduced while mesenchymal and endothelial markers were elevated (**Fig. S3D**). These results suggest that the initial cellular composition in the bone marrow microenvironment and its alteration by combination therapy might contribute to better outcomes in HMA-R/R MDS patients.

### Specific cellular composition and dynamics mark STS and LTS bone marrow

Given that tumor cell-extrinsic factors in the bone marrow could be shaping HMA-R/R MDS patient survival, we were motivated to resolve the CD34+ cellular subtypes and immune cell composition in the bone marrow using single-cell sequencing. Therefore, we performed CITE-seq^31,32^ on available bone marrow samples from three STS patients and 10 LTS patients, before treatment and after two treatment cycles (**Table S2**). This method enabled us to quantify whole transcriptomes and the abundance of ∼150-200 cell surface proteins at single-cell resolution (see **Methods, Table S3-S4**). After filtering for high-quality cells, we used Seurat^33,34^ and the multimodal information to integrate and cluster cells into 30 distinct HSPC, stromal, or immune subtypes in the bone marrow microenvironment (verified by cell type-specific marker expression; **Fig. S4-6**)^34–36^. In total, we generated an atlas of 61,487 cells that reflect the bone marrow microenvironment heterogeneity in the patients before and after combination therapy (**Fig. S7A**).

Next, we compared the pre-treatment bone marrow cellular composition between STS and LTS to determine whether specific HSPC or immune cell subtypes were disproportionately represented at baseline. Surprisingly, the hematopoietic stem cell (HSC), lymphoid-primed multipotent progenitor (LMPP), and granulocyte macrophage progenitor (GMP) fractions were higher in the bone marrow of LTS relative to STS (**Fig. 2A**). These stem or progenitor cell populations are known to contain the malignant and disease-initiating cells. We also found that naïve CD4 and CD4 T central memory cells were significantly enriched in the STS bone marrow, which could reflect the dysfunction or paucity of dendritic cells to properly transactivate CD4 T cell for differentiation. Indeed, LTS had more antigen-presenting cells (myeloid dendritic cells and plasmacytoid dendritic cells) in the bone marrow than STS, which coincided with a significant expansion of effector lymphocytes (natural killer, gamma-delta T, CD4 and CD8 T cells) and reduction of HSPCs (HSC, LMPP, GMP) after combination therapy (**Fig. 2B**). In the bone marrow of STS, there were decreased CD4 and CD8 T cell numbers but increased HSC and LMPP cells after combination therapy, demonstrating failure to reinvigorate the bone marrow microenvironment immunogenicity and the subsequent, immediate disease progression. Thus, HMA-R/R MDS patients have distinct bone marrow microenvironment cellular compositions at baseline and after combination therapy that reflect MDS progression or resolution.

**Figure 2.**
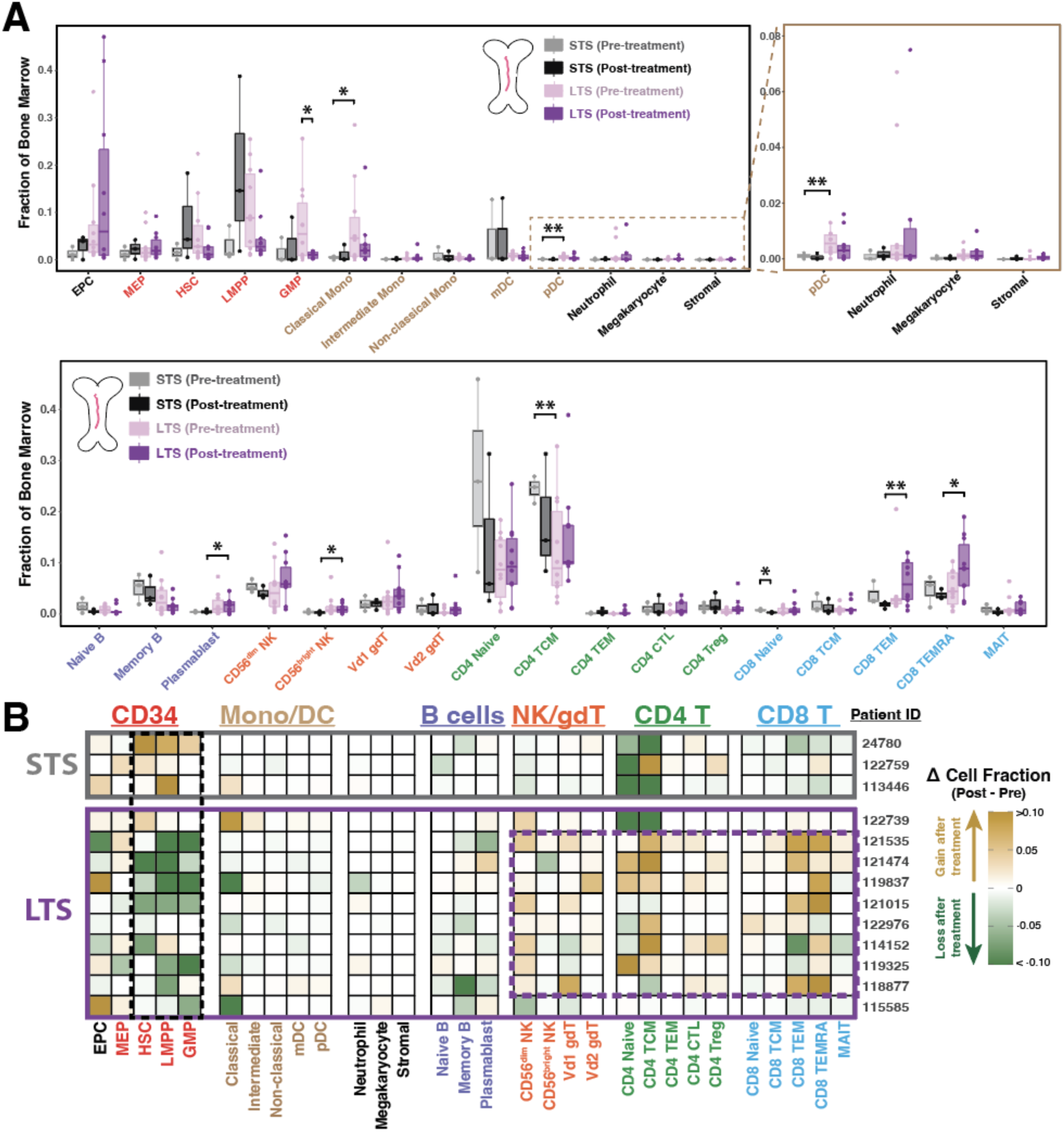
Distinct immune compositions and changes detected in the bone marrows of HMA-R/R MDS patients treated with combination therapy. **A.** The distribution of cell subtype fractions detected in the bone marrow of STS or LTS patients before and after combination therapy. Brown box presents magnified boxplots of bone marrow cellular fractions. (Two-tailed Welch’s t-test: * denotes p-value < 0.05, ** denotes p-value <0.01). **B.** Heatmap reflecting the changes in bone marrow cellular fractions for each patient after combination therapy. Erythroid precursor cells, EPC; Megakaryocyte erythroid progenitor, MEP; Hematopoietic Stem Cell, HSC; Lymphoid primed multipotent progenitor, LMPP; Granulocyte-monocyte progenitor, GMP; Myeloid dendritic cell, mDC; Plasmacytoid dendritic cell, pDC; Natural Killer, NK; Gamma-delta T, gdT; T central memory, TCM; T effector memory, TEM; Terminally differentiated effector memory, TEMRA; cytotoxic, CTL; regulatory T, Treg; Mucosal-associated invariant T, MAIT.

### STS bone marrow demonstrates elevated inflammation and senescence-like signatures in HSPCs

Considering the rapid disease progression in STS, we searched for gene pathway signatures that might explain the early bone marrow microenvironment dynamics that contributed to therapy failure. We used pseudo-bulk comparisons (**Methods**, DESeq2^37^) for GSEA to minimize the impact of patient-to-patient variation or imbalanced cell numbers across groups^38^. Any cell subtypes with less than 10 cells detected in a sample were excluded from pseudo-bulk analysis. Indeed, pseudo-bulk analysis of STS-CD34+ cells (combined MEP, HSC, LMPP, and GMP cells) reproduced the inflammation and oncogenic pathway signatures that were enriched in bulk bone marrow CD34+ cells from STS (**Fig. S7B**). Interestingly, these pseudo-bulk STS-CD34+ cells showed treatment-induced interferon gamma/alpha activation, but lacked the proliferation potential as seen in the LTS-CD34+ cells. In contrast to bulk GSEA results, we found that STS-CD34+ cells (specifically the LMPP subtype) were enriched in the EMT pathway, but this enrichment was decoupled from collagen-related pathways (**Fig. S7C**). This discordance might arise because mesenchymal stromal cells (CXCL12-expressing stromal cells^39,40^) had weak CD34 gene and protein expression, and were excluded from the analysis. CITE-seq was not able to capture a sufficient number of mesenchymal stromal cells from bone marrow aspirates, thus further work is needed to evaluate the contributions of mesenchymal stromal cells to clinical responses in these MDS patients.

We then expanded the hallmark GSEA comparison to the specific cell subtypes, and discovered that the pseudo-bulk CD34+ GSEA hallmark signatures were mirrored in the HSPC subpopulations (pseudo-bulk comparisons of MEP, HSC, LMPP) in STS bone marrow (**Fig. 3A**). These results suggest that high inflammatory response in STS-HSPCs might be tied to slow proliferation. Next, to search for candidate genes that provide prognostic value for combination therapy failure, we performed single-cell differential analysis (**Methods,** Wilcoxon rank sum test^34^). We found that STS-HSPCs had increased expression of *CDKN2C, CDKN2D, CXCL2, CXCL3,* and *CXCL8* (**Fig. 3B**). *CDKN2C* (p18) and *CDKN2D* (p19) belong to the cyclin-dependent kinase inhibitor family, which control cell proliferation and promote cellular senescence^41–43^. *CXCL2, CXCL3,* and *CXCL8* (IL-8) are proinflammatory senescence-associated secretory phenotype (SASP) cytokines, which are secreted by senescent cells to create a highly inflamed microenvironment with significant immune suppression and dysfunction^44,45^. CDKN and CXCL gene expression was elevated in most STS-HPSCs, especially in LMPPs (**Fig. S8A-C**). Pseudo-bulk GSEA of senescence-associated pathway (“SenMayo”^44^) also confirmed that STS-LMPPs have increased senescent-like signatures that were not resolved by combination therapy (**Fig. 3C**). These senescence-like signatures were also enriched in bulk transcriptomes of bone marrow STS-CD34+ cells at baseline (**Fig. S8D**). Indeed, ligand-receptor inference as shown using CellChat^46^ revealed that tumor necrosis factor (TNF) and IL-1 signaling interactions (which associate with chronic inflammation and senescence^47^) were specifically detected in the STS-HSPC subpopulations, but not in LTS-HSPCs (**Fig. S9**). In a previous study, *ITGA5* up-regulation and cellular quiescence was associated with initial HMA therapy failure in MDS patients^48^. In our small patient cohort, we only detected *ITGA5* up-regulation in STS-HSCs, but not in other HSPC subpopulations (**Fig. S8E**). These data suggest that high inflammation and the presence of senescent-like cells in the bone marrow microenvironment might be a prognostic biomarker for patients who will not respond to combination therapy.

**Figure 3.**
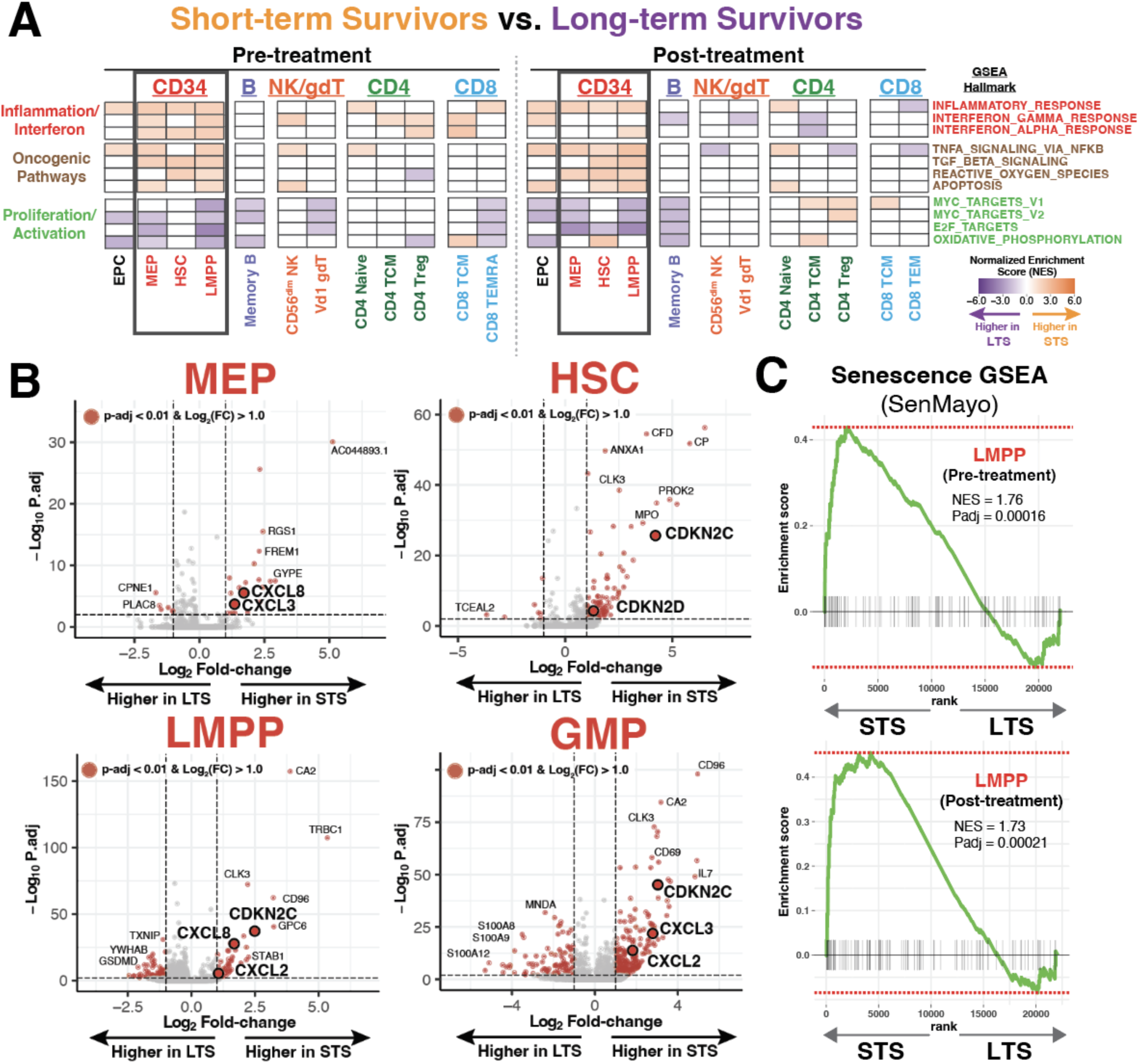
Enhanced inflammation and senescent-like signatures in bone marrow HPSCs of short survivors. **A.** Pseudo-bulk GSEA of hallmark pathways comparing STS and LTS bone marrow cell subtypes across treatment timepoints (adjusted p-value < 0.05). **B.** Volcano plots illustrating single-cell differential gene expression analysis comparing STS- and LTS-HSPCs at study entry. **C.** GSEA of the senescence pathway, “SenMayo”, detected in LMPP cells of short-term survivors.

### Combination therapy eliminates immunosuppressive monocytes in LTS bone marrow

LTS-bone marrow microenvironment cellular dynamics revealed that combination therapy expanded lymphocytes and eliminated HSPCs, which potentially contributed to the better outcome in the LTS patients. Pseudo-bulk GSEA comparing treatment timepoints unveiled robust interferon pathway activations in most of the LTS bone marrow cells, which were remarkably absent in the STS bone marrow cells (**Fig. 4A & S10**). Thus, combination therapy likely reinvigorated effector immune cell cytotoxic functions in LTS bone marrow microenvironment that could contribute to the elimination of the malignant myeloid cells/blasts in the bone marrow.

**Figure 4.**
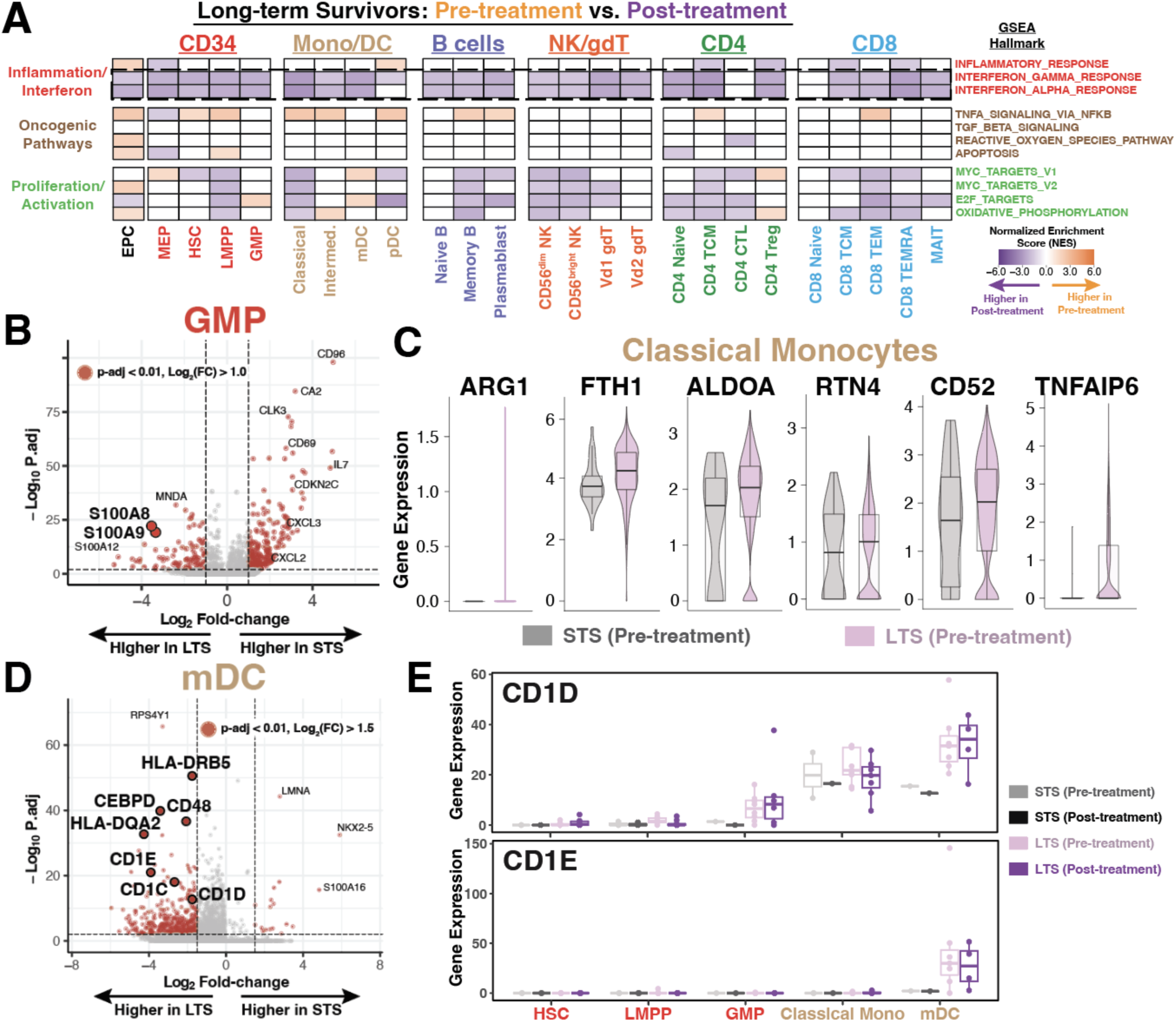
LTS bone marrow microenvironments are enriched with S100A8/9-expressing myeloid cells and CD1-expressing mDCs. **A.** Pseudo-bulk GSEA of hallmark pathways comparing pre-treatment and post-treatment samples of LTS cell subtypes (adjusted p-value < 0.05). **B.** Single-cell differential gene expression analysis comparing STS- and LTS-GMPs at pre-treatment timepoint. **C.** Distribution of MDSC-marker gene expression in classical monocytes of STS or LTS at pre-treatment timepoint. **D.** Single-cell differential gene expression analysis comparing STS- and LTS-mDCs at pre-treatment timepoint. **E.** Pseudo-bulk gene expression profiles of differentially expressed *CD1D* and *CD1E* in HSPCs and myeloid cells. Myeloid-derived suppressor cells, MDSC.

Previously, we noticed that LTS bone marrow had more GMP and myeloid cells relative to STS bone marrow (**Fig. 2A**). The transcriptomic comparisons revealed that *S100A8* and *S100A9* alarmin genes were preferentially expressed in GMPs and classical monocytes from the LTS bone marrow (**Fig. 4B & S11A,B**). While alarmin genes are often associated with inflammation, *S100A8/9* secretion can trigger PD-1 expression in HSPCs^49,50^ (as supported by **Fig. S12**) and recruit myeloid-derived suppressor cells (MDSCs) to the bone marrow, thus transforming the bone marrow microenvironment into an immunosuppressive and tumor-promoting microenvironment^51–53^. MDSCs propagate alarmin signaling pathways by becoming an additional source of *S100A8/9* production in the bone marrow^54^. It is therefore possible that MDSC recruitment is reflected in the increased myeloid presence in LTS bone marrow. Indeed, we detected elevated monocytic MDSC-marker gene^55^ expression in the LTS-classical monocytes, suggesting higher prevalence of MDSCs in the LTS bone marrow than STS bone marrow at baseline (**Fig. 4C**). Combination therapy drastically reduced GMPs and classical monocytes in LTS bone marrow (**Fig. 2B**), which likely reflect the therapeutic elimination of MDSCs in the bone marrow. Therefore, HMA-R/R patients with high MDSC infiltration in their bone marrow might benefit from combination therapy due to the ICI-mediated clearance of MDSC immunosuppression and disease burden.

### Non-canonical, dendritic cell antigen presentation is associated with patient survival

Interactions between dendritic cells and T cells are critical for tumor clearance^56^. The STS bone marrow at baseline was mostly comprised of naïve CD4 or CD4 TCM cells (**Fig. 2B**), implying that STS could have compromised dendritic cell function relative to LTS. Indeed, LTS bone marrow had greater clonal expansion of CD4 and CD8 T cells (as estimated by the T cell receptor (TCR) profiling tool TRUST4^57^; **Fig. S11B,C**) after combination therapy, which coincided with the increased myeloid dendritic cells (mDCs) and plasmacytoid dendritic cells (pDCs) as detected by CITE-seq. Traditional antigen-presenting cells are primarily attributed to mDCs, while pDCs are characterized by the production of interferon gamma in the bone marrow microenvironment^58^. We therefore profiled mDCs to gain mechanistic insights into superior T cell activation in long survivor bone marrow microenvironments.

At baseline, LTS-mDCs up-regulated human leukocyte antigen (HLA) class II molecules and CD48, both of which are critical for CD4 T or NK cell activation^59^ (**Fig. 4D**). There was also increased expression of *CEBPD*, which is often associated with inflammation and *S100A8/9* induction in myeloid cells^60^. A subset of CD1 glycoprotein family genes (*CD1D* and *CD1E*) were also preferentially expressed in LTS-mDCs (**Fig. 4E**). Interestingly, *CD1D* gene and protein expression was also elevated in LTS-LMPP and LTS-GMP cells (**Fig. S11E**). Lipid antigens are presented by CD1D molecules, and ψ-δ TCRs on Vd1 gdT cells recognize these lipid antigens to induce effector function^61,62^. There was a modest expansion of gdT cells in LTS bone marrow after combination therapy (**Fig. 2C**). Therefore, we speculate that LTS-mDCs were already primed for peptide and lipid antigen presentation before treatment, but were likely suppressed by MDSCs. In conclusion, we suggest that the addition of ICI resolved the MDSC-mediated immunosuppression in the LTS bone marrow microenvironment, thus stimulating cytotoxic NK, gdT, CD4, and CD8 T cells (via DC interaction), and ultimately reducing tumor burden in patients.

### Evaluating peripheral blood samples as surrogates for bone marrow microenvironment dynamics

Bone marrow biopsies and peripheral blood samples are often collected from patients with hematological malignancies for correlative and follow-up studies, but it is not clear if PBMCs accurately reflect bone marrow microenvironment dynamics in MDS patients who receive combination therapy. In our previous work, we extensively profiled NK and CD8 T cell dynamics in peripheral blood using flow cytometry-based immunophenotyping, but did not detect a prognostic immune signature that segregated STS and LTS patients^19^. We hence generated and analyzed CITE-seq data from available PBMC samples (**Fig. S7A**) to compare and identify prognostic molecular signatures associated with patient survival after combination therapy. Before treatment, immune cell compositions in STS- and LTS-PBMCs were similar to what we observed in their corresponding bone marrows, with the exception of HSPCs (**Fig. 5A**). The abundance of conventional immune checkpoints (PD-1, TIGIT, LAG-3, and TIM-3) on PBMC immune cells was also similar to their corresponding bone marrow immune cells (**Fig. S13**). Thus, PBMCs can accurately reflect bone marrow cells’ immune checkpoint expression. After combination therapy, however, PBMC immune cell composition changes did not reflect bone marrow microenvironment dynamics (**Fig. 5B**). Thus, acquiring bone marrow biopsies after treatment could be critical for evaluating therapy efficacy in MDS patients after combination therapy.

**Figure 5.**
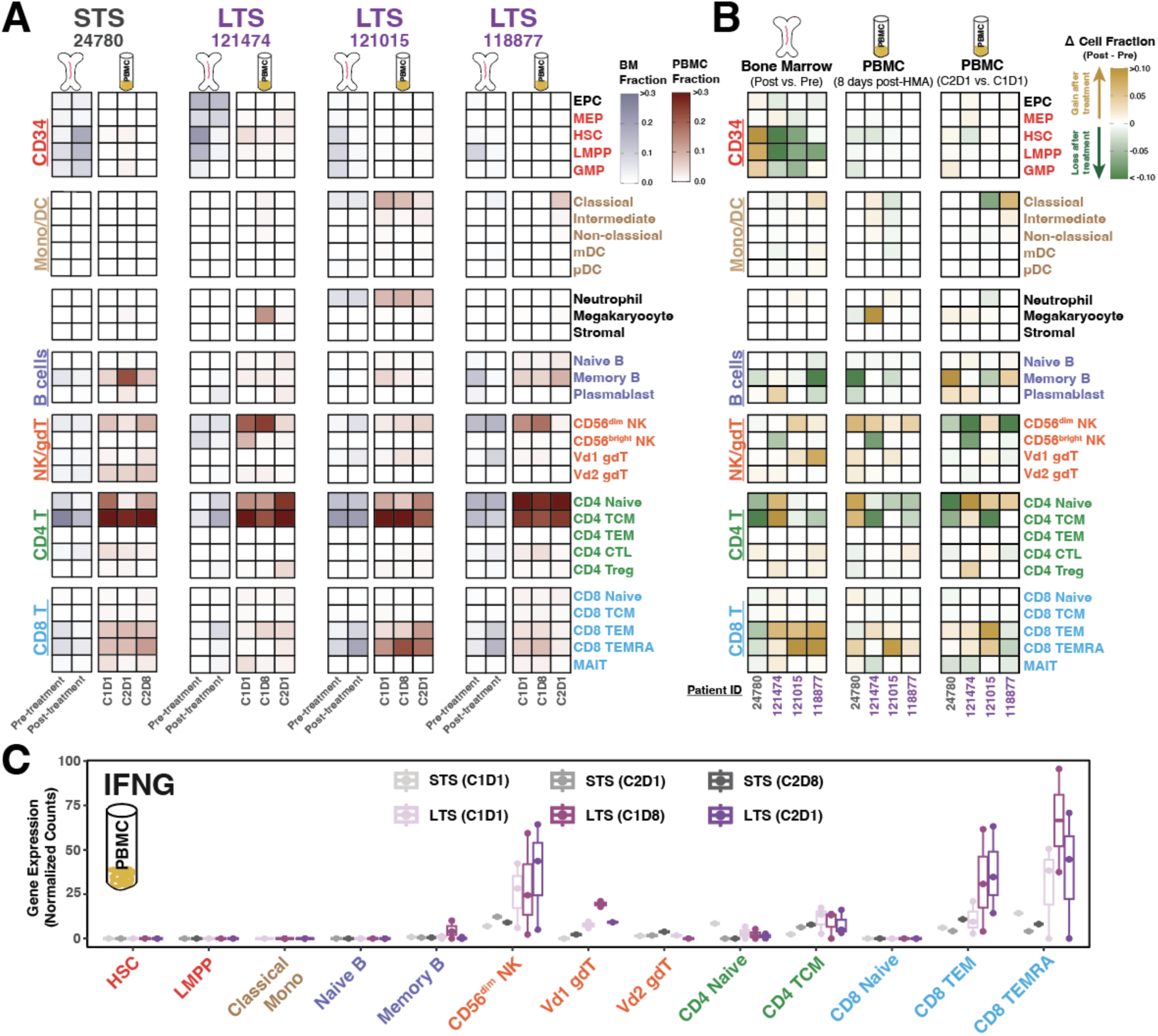
CITE-seq profiling of PBMCs from HMA-R/R MDS patients treated with combination therapy. **A.** Heatmaps illustrating the fraction of cell subtypes detected in the bone marrow or peripheral blood mononuclear cell (PBMC) samples of STS and LTS patients across treatment timepoints. **B.** Heatmap reflecting the change in cellular fractions in the bone marrow or PBMC samples for select MDS patients after combination therapy. **C.** Pseudo-bulk expression distribution of *IFNG* in PBMC immune subtypes.

We also asked if differential activation of treatment-induced viral mimicry pathways in PBMCs was associated with patient survival, but we did not detect any correlation between viral mimicry activation and survival (**Fig. S14A,B**). Instead, we found that basal and treatment-induced *IFNG* (interferon gamma) up-regulation in effector lymphocytes segregated STS and LTS (**Fig. 5C**). We also found that senescent-like or alarmin signatures discovered in the bone marrow microenvironment were also present in PBMC samples; senescent-associated genes were elevated in peripheral CD34+ cells of STS-PBMC, and high *S100A8/9* expression was detected in the peripheral monocytes of LTS (**Fig. S14C,D**). Thus, PBMCs can reflect baseline bone marrow microenvironment immune composition, but not treatment-induced bone marrow microenvironment dynamics, and IFNG, senescent-like, or alarmin signatures in PMBCs might be useful for stratifying HMA-R/R MDS patients for combination therapy.

## Discussion

There is a significant clinical need to design new treatments for HMA-R/R MDS patients. We recently reported the results of a Phase I/II clinical trial that combined guadecitabine (HMA) with atezolizumab (anti-PD-L1) as a way to potentially reverse HMA resistance mediated by immune checkpoint up-regulation^19^. The overall response rate at 6 months was 30%, and the median overall survival for patients receiving combination therapy was extended to 15.1 months, which prompted us to identify potential mechanisms that contributed to prolonged survival. Previously, we highlighted the association between longer patient survival and the detection of *ASXL1* mutations in both myeloid and T cells^19^. While we have shown that the deletion of *Asxl1* in CD8 T cells enhances their self-renewal capacity and the durability of their responsiveness to ICIs^63^, CITE-seq was not able to reliably capture *ASXL1* expression in various immune cell types to discriminate ASXL1-mutated cells for further interrogation in this study. Instead, we present here the transcriptional signatures of bone marrow cells and bone marrow microenvironment dynamics (**Fig. 6**) that associate with improved survival in HMA-R/R MDS patients treated with combination therapy.

**Figure 6.**
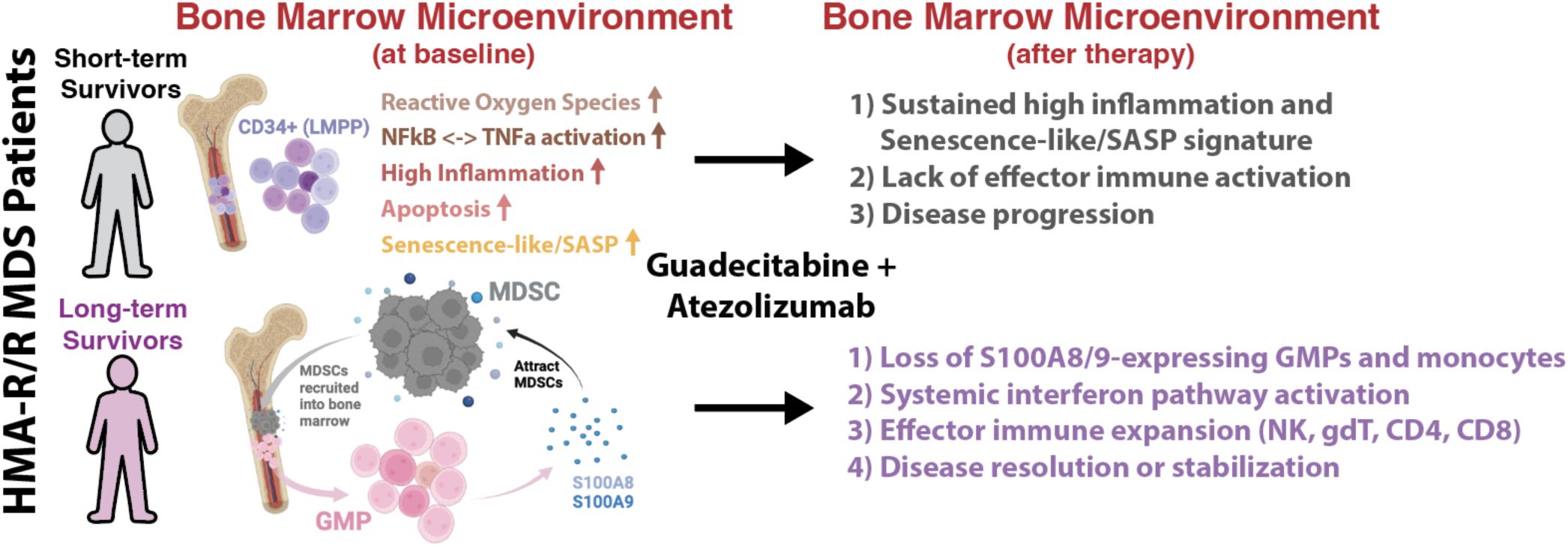
Two bone marrow microenvironments segregate STS and LTS patients in HMA-R/R MDS patients. Simplified schematic illustrating two bone marrow microenvironments, senescence-associated inflammation and MDSC-mediated immunosuppression, that segregate STS or LTS patients after combination therapy respectively. Tumor necrosis factor alpha, TNFa; nuclear factor kappa B, NFkB; Senescence-associated secretory phenotype, SASP. Illustrations were created with BioRender.

Bulk bone marrow CD34+ transcriptome comparisons revealed that combination therapy induced interferon responses in patient-derived CD34+ cells, which was decoupled from the transcriptional reactivation of TEs. Instead, LTS-CD34+ cells were enriched for collagen-containing ECM and EMT pathways after combination therapy. These data suggested that extrinsic immunomodulatory factors or CD34+ cell type composition (which include HSPC, MSC, and endothelial cells) in the bone marrow microenvironment could shape clinical response. We therefore generated a single-cell atlas of the bone marrow microenvironment from a subset of HMA-R/R MDS patients. While we were not able to capture adequate number of MSCs in the CITE-seq data for analysis, we corroborated the increased interferon and effector immune response after combination therapy in LTS bone marrow microenvironment. In contrast, the STS-bone marrow microenvironment at baseline was marked with an overabundance of naïve CD4 T and CD4 TCM cells, which suggest CD4 T cell differentiation defects. Further gene expression and pathway analysis suggested that high inflammation, elevated TNFα signaling (via NFkB in HSPCs), and senescent-like CD34+ cells could have produced an immunosuppressive environment in the STS bone marrow, which could not be ameliorated by combination therapy. Senescence-like signatures have been previously tied to therapy failure in heme malignancy^64^, and therapeutic strategies targeting senescent cells, such as senolytics, could be a better therapeutic alternative in these particular patients^65^.

The LTS bone marrow microenvironment was enriched with myeloid cells, and LTS-GMPs and LTS-classical monocytes showed elevated expression of *S100A8/9* and MDSC-marker genes. These results suggest that pre-existing or acquired immunosuppressive myeloid cells blocked the therapeutic efficacy of the initial HMA treatment. However, adding atezolizumab (an immune checkpoint inhibitor) reconciled MDSC-mediated immunosuppression, thus reinvigorating global effector immune functions and reducing malignant cancer cells and MDSCs (**Fig. S15A**). While canonical CD4 T cell – CD8 T cell – dendritic cell interactions are critical for immune-mediated cancer eradication^56^, we observed that myeloid dendritic cells in LTS bone marrow were also primed to present lipid antigens through CD1D and CD1E molecules, which can be recognized by Vd1 gdT cells to induce anti-tumor effector consequences and reduce disease burden^62^ (**Fig. S15B**). These data suggest that ICIs can also promote noncanonical antigen presentation and T cell interactions, which is currently underappreciated in the context of MDS correlative studies.

In conclusion, we have identified two signatures in the bone marrow microenvironment that associate with survival in HMA-R/R MDS patients treated with an HMA (guadecitabine) and ICI (atezolizumab) combination therapy. We suggest that these bone marrow biomarkers might be useful for stratifying and selecting patients who will benefit from the combination therapy, however this will need to be validated in a larger study. Discerning which molecular markers or mechanisms predict ineffective or bad prognosis after combination treatment will also assist in prioritizing alternative therapeutic interventions, and optimizing care for at-risk populations. We hope that this study can be a valuable resource, and help shape the next iteration of clinical trials in this vulnerable patient population for which there are limited alternative therapies.

## Methods

### Bulk totalRNA-seq of bone marrow CD34+ cells

Bone marrow mononuclear cells (BMMCs) were isolated from patients as described in our previous work^19^, and were cryopreserved in liquid nitrogen in the VAI Biorepository Core. BMMCs were quickly thawed and resuspended by slowly adding thawing medium (DPBS + 5% FBS (Sigma-Aldrich) + 60U/ml DNase I + 5mM MgCl2) in stepwise fashion. CD34+ cells were separated through magnetic purification (Miltenyi CD34 UltraPure) and total RNA was extracted with Trizol (Thermo Fisher Scientifc) – isopropanol method. The VAI Genomics Core generated bulk totalRNA-seq libraries using a QIAseq FastSelect (Qiagen) and KAPA RNA HyperPrep Kit (Roche) and sequenced the libraries using the Illumina NovaSeq6000 platform. Healthy donor bone marrow CD34+ total RNA-seq libraries were generated in a separate study and deidentified before analysis. Total RNA-seq libraries were adapter trimmed (TrimGalore, https://zenodo.org/records/7598955), filtered *in silico* for ribosomal RNA^66^, down-sampled to 35 million paired reads to normalize library size, and aligned using STAR^67^. Differential gene expression and pathway analysis was performed using DESeq2^37^ (v.1.42.0) and fgsea^68^ (v.1.28.0). TE transcript quantification was performed using featureCounts^69^ as previously described^70^.

### Bone marrow and PBMC CITE-seq

BMMC and PBMC cryopreserved vials were thawed and resuspended as mentioned above. Five to six samples were incubated with TotalSeq-C Hashtag antibodies (BioLegend) and sorted for live cells using flow cytometry. We pooled samples and labeled cells with TotalSeq-C antibody cocktail (BioLegend) following the manufacturer’s instructions. Single-cell gene and surface protein libraries were constructed with 10x Chromium Next GEM 5’ GEM Kit v1.1 (10x Genomics) and sequenced on the Illumina NovaSeq6000 platform. Gene and protein count matrices were generated with Cell Ranger (v4.0.0) and processed with CellBender^71^ (v0.3.0) to remove ambient or background noise. Gene and protein count matrices were normalized with SCTransform and integrated using weighted-nearest neighbor analysis workflow with Seurat^33^ (v5.0.1). Cell cluster identification was guided by Azimuth^34^ (v0.5.0) followed by manual curation and verification with established cell-type marker genes. The FindMarkers function (Wilcoxon ran sum test) was used for single-cell differential expression testing. Pseudo-bulk count matrices for each cell subtypes were generated using SingleCellExperiment^72^ (v1.24.0) and GSEA analysis was performed with DESeq2 and fgsea packages. Any cell subtypes with <10 cells detected in a sample were excluded from pseudo-bulk analysis. We used TRUST4^57^ (v.1.0.13) for T cell receptor and clonal analysis, and CellChat^46^ (v2.1.2) for ligand-receptor analysis. All analyses were performed in R v4.3.0 (RStudio 2024.02.999). Sequencing data is being deposited to Gene Expression Omnibus and will be released upon final publication.

## Acknowledgements

The authors thank VAI-SU2C Epigenetics Team members and Jones lab members for providing valuable scientific feedback for this work; M. Tang for valuable resources and tutorials for single cell analysis; D. Chandler and D. Brass for editorial assistance with the manuscript. We also thank all the members of the VAI Genomics Core (RRID:SCR_022913), VAI Flow Cytometry Core (RRID:SCR_022685), VAI Bioinformatics and Biostatistics Core (RRID:SCR_024762), and VAI Pathology & Biorepository Core (RRID:SCR_022912) for their help with sample storage, processing, and sequencing library construction. We are especially grateful to the patients who provided valuable samples for this study. Research funding was provided by Van Andel Institute through the Van Andel Institute – Stand Up To Cancer Epigenetics Dream Team. This work was also supported by funding from National Institutes of Health, National Cancer Institute (K99CA286742 to H.J.J, R03CA290259 to T.J.T.J. and R35CA209859 to P.A.J.).

## Authorship Contribution

Contribution: H.J.J. and P.A.J. conceptualized the project; H.J.J., R.S.B., M.A., and R.S. designed experiments; H.J.J., G.U., A.D.O., H.J.K., S.A.N., A.V.N., H.L., M.W., and K.B. performed experiments; H.J.J. analyzed data with assistance from Z.H.R., S.A.G., B.J.K., C.C.Z., B.A.Y., J.J.I., M.J.T., S.B.B., M.R.B., T.J.T.J., C.L.O., and K.G.; H.J.J. and P.A.J. wrote the manuscript; and all authors edited the manuscript.

## Supplemental Tables

**Supplemental Table 1 – Bulk CD34+ RNA-seq sample information**

**Supplemental Table 2 – CITE-seq sample information**

**Supplemental Table 3 – TotalSeq-C Cocktail 1 Barcodes**

**Supplemental Table 4 – TotalSeq-C Cocktail 2 Barcodes**

---

**Supplemental Figure 1.**
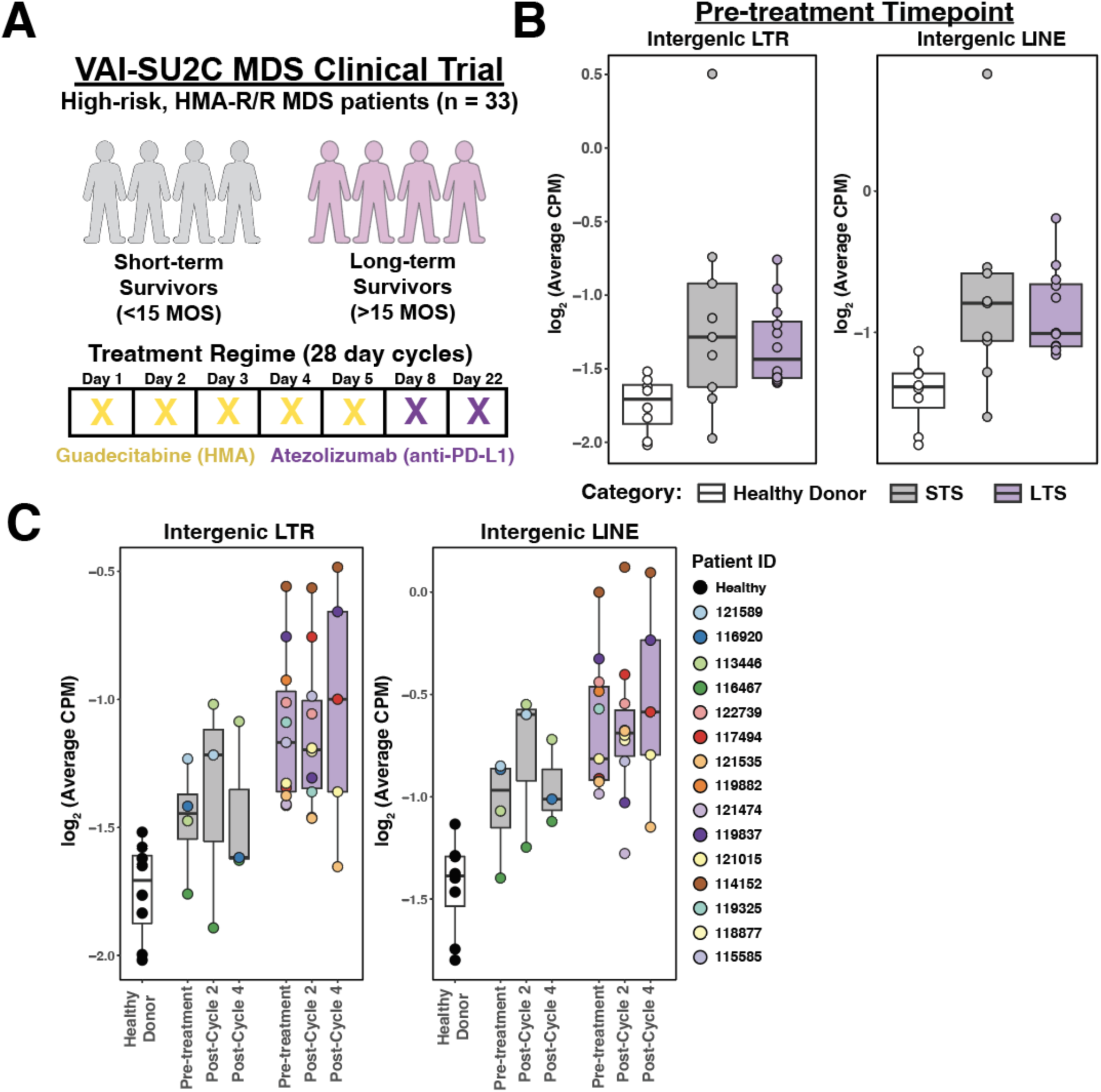
Quantification of intergenic TE transcripts detected in bulk CD34+ cells from bone marrow of HMA-R/R MDS patients. **A.** Treatment and sample collection overview of VAI-SU2C MDS clinical trial combining guadecitabine and atezolizumab in HMA-R/R MDS patient cohort. **B.** Comparison of intergenic LTR (left) or intergenic LINE (right) transcript abundance detected in CD34+ cells isolated from healthy donor, STS, and LTS bone marrow at pre-treatment timepoint. **C.** Quantification of intergenic LTR (left) or intergenic LINE (right) transcripts in healthy donor CD34+ or patient-derived CD34+ cells from bone marrow across treatment cycles after combination therapy. Months overall survival, MOS.

**Supplemental Figure 2.**
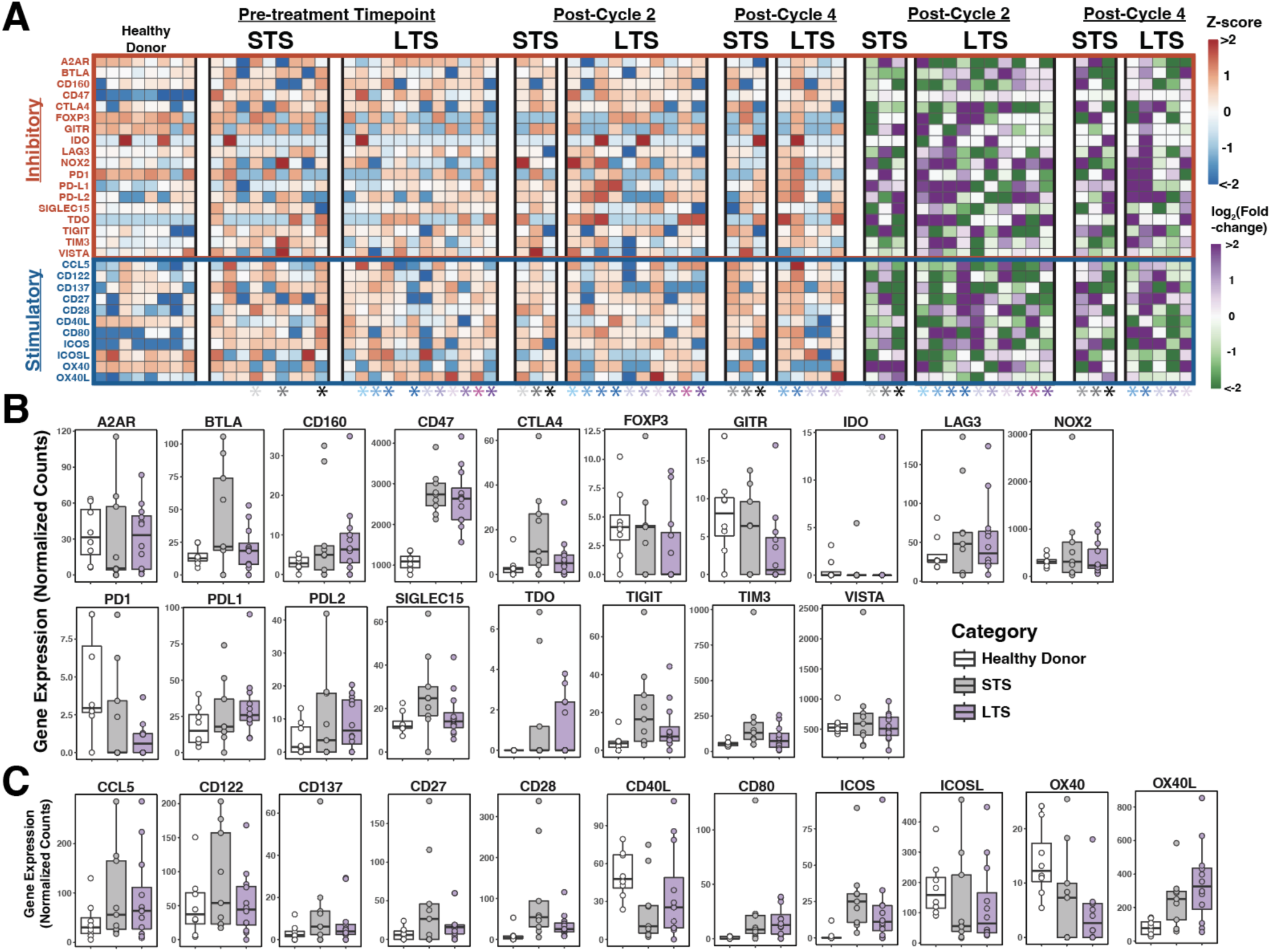
Immune checkpoint gene expression dynamics in healthy or patient-derived bone marrow CD34+ cells. **A.** Heatmaps representing immune checkpoint gene expression (left) or fold-change (right) after combination therapy in healthy donor CD34+ or patient-derived bone marrow CD34+ cells. **B-C.** Distribution of inhibitory (**B**) or stimulatory (**C**) immune checkpoint gene expressions in CD34+ cells isolated from healthy donor, STS, and LTS bone marrow at pre-treatment timepoint. Colored asterisks represent patient ID.

**Supplemental Figure 3.**
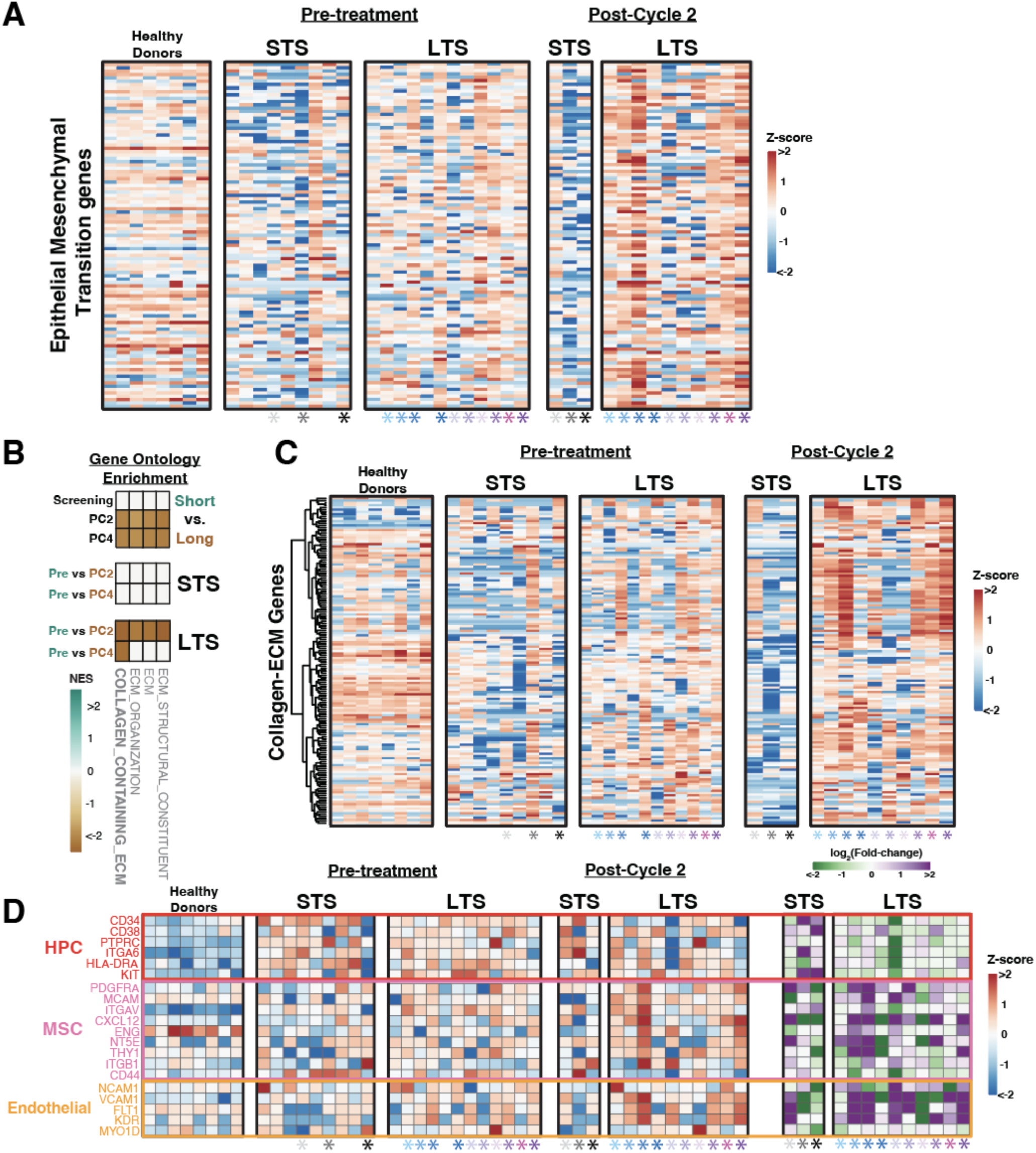
Therapy-induced collagen-ECM pathway signatures and CD34+ cell type heterogeneity enriched in long survivors’ bone marrow microenvironment. **A.** Heatmaps representing expression of Epithelial Mesenchymal Transition (EMT) pathway genes after combination therapy in healthy donor CD34+ or patient-derived CD34+ cells from the bone marrow. **B.** Gene Set Enrichment Analysis using gene ontology (Cellular Component) comparing STS or LTS CD34+ cells across treatment timepoints (adjusted p-value < 0.01). **C.** Expression of collagen-containing-ECM pathway genes in healthy donor CD34+ or patient-derived CD34+ cells from bone marrow before and after combination therapy. **D.** Heatmaps representing CD34+ subtype marker gene expression (left) or fold-change (right) after combination therapy in healthy or patient-derived bone marrow CD34+ cells. Colored asterisks represent various patient IDs.

**Supplemental Figure 4.**
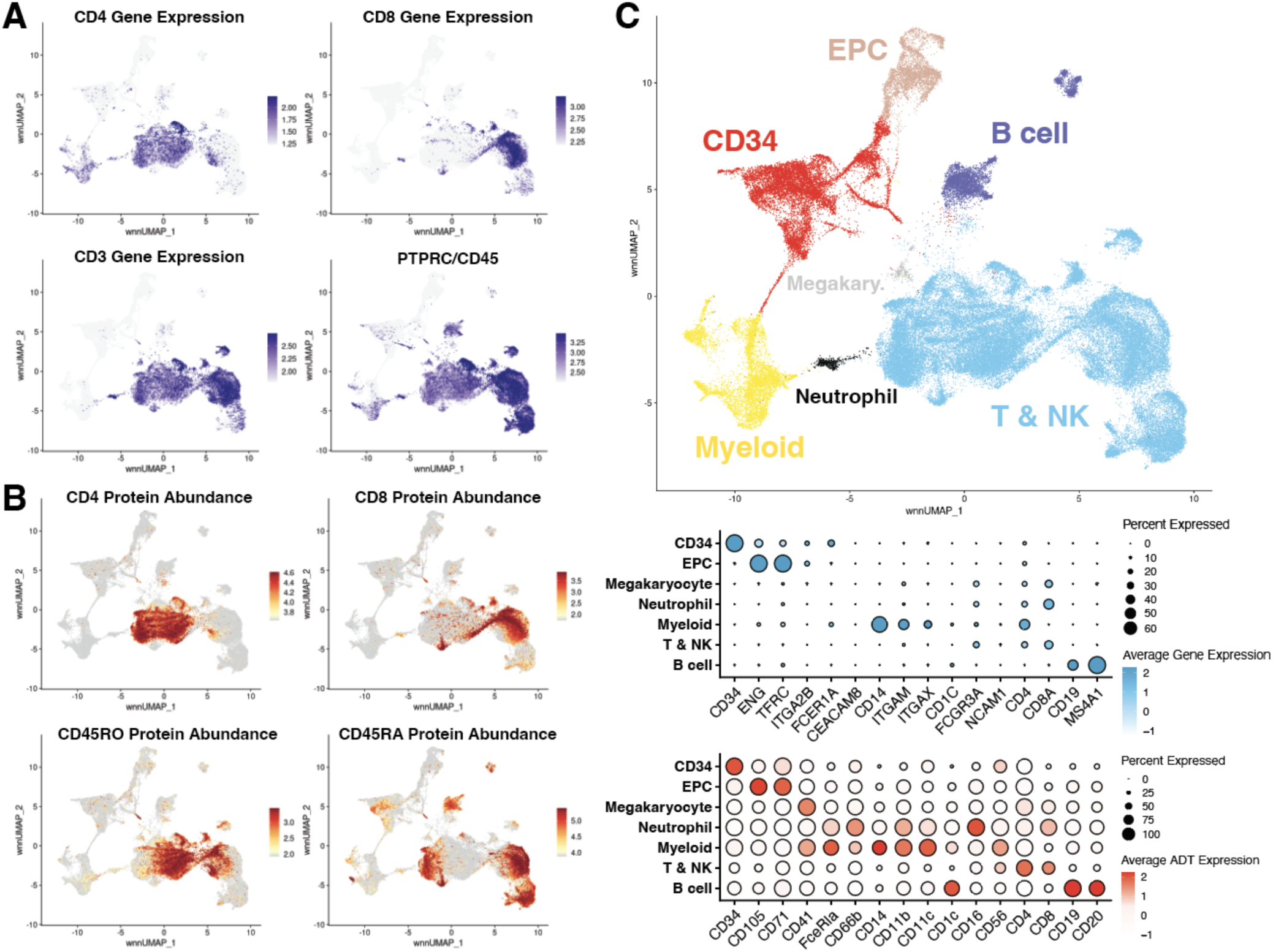
Broad cell type annotation of bone marrow microenvironment using multimodal information from CITE-seq. **A-B.** Gene expression (**A**) or protein abundance (**B**) of broad cell type markers (CD4, CD8, CD45RO, CD45RA) quantified in each single cell detected in the bone marrow microenvironment, which are projected on weighted nearest neighbor-based UMAP (wnnUMAP). **C.** The wnnUMAP projection of the 7 broad cell types identified in the bone marrow microenvironment of MDS patients (top) and gene and protein expression dot plots (bottom) of marker genes for each broad cell type clusters.

**Supplemental Figure 5.**
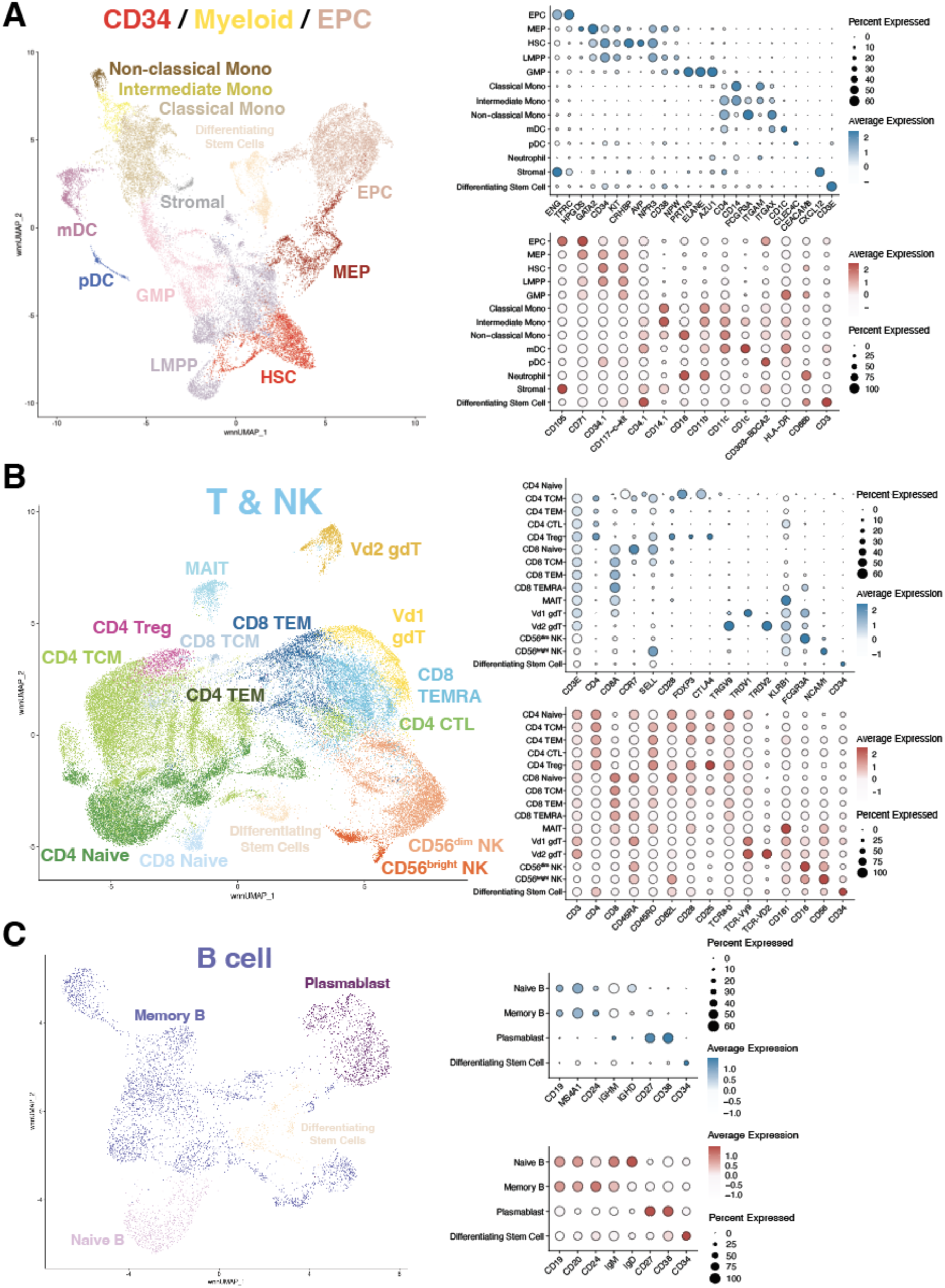
Specific cell subtype annotation after re-clustering broad cell type clusters of CITE-seq data. **A-C.** The wnnUMAP projection of the various cell subtypes identified in CD34-myeloid-EPC (**A**), T & NK (**B**), and B cell (**C**) groups detected in the bone marrow microenvironment of MDS patients (left) and gene and protein expression dot plots (right) of marker genes for each specific cell subtype clusters.

**Supplemental Figure 6.**
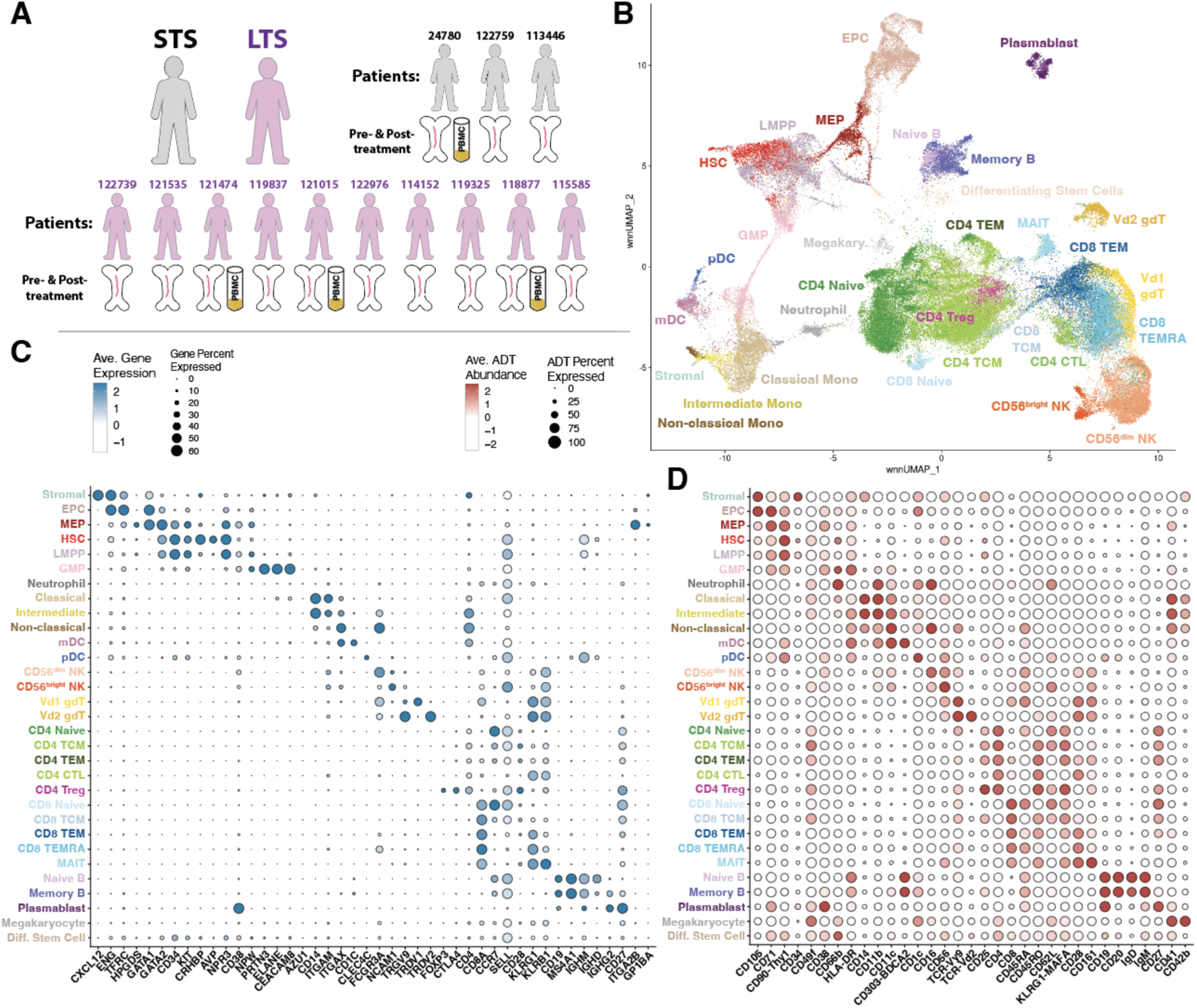
Marker gene validation of cell subtypes detected in bone marrow microenvironment and PBMCs of MDS patients. **A.** Schematic illustrating the types of patient samples processed. **B.** Weighted nearest neighbor-based UMAP (wnnUMAP) projection of the 30 cell subtypes identified in the bone marrow microenvironment of MDS patients. **C-D.** Dot plot illustrating gene expression (**C**) or protein abundance (**D**) of various cell subtype-specific markers in 30 specific cell subtype clusters detected in the CITE-seq data.

**Supplemental Figure 7.**
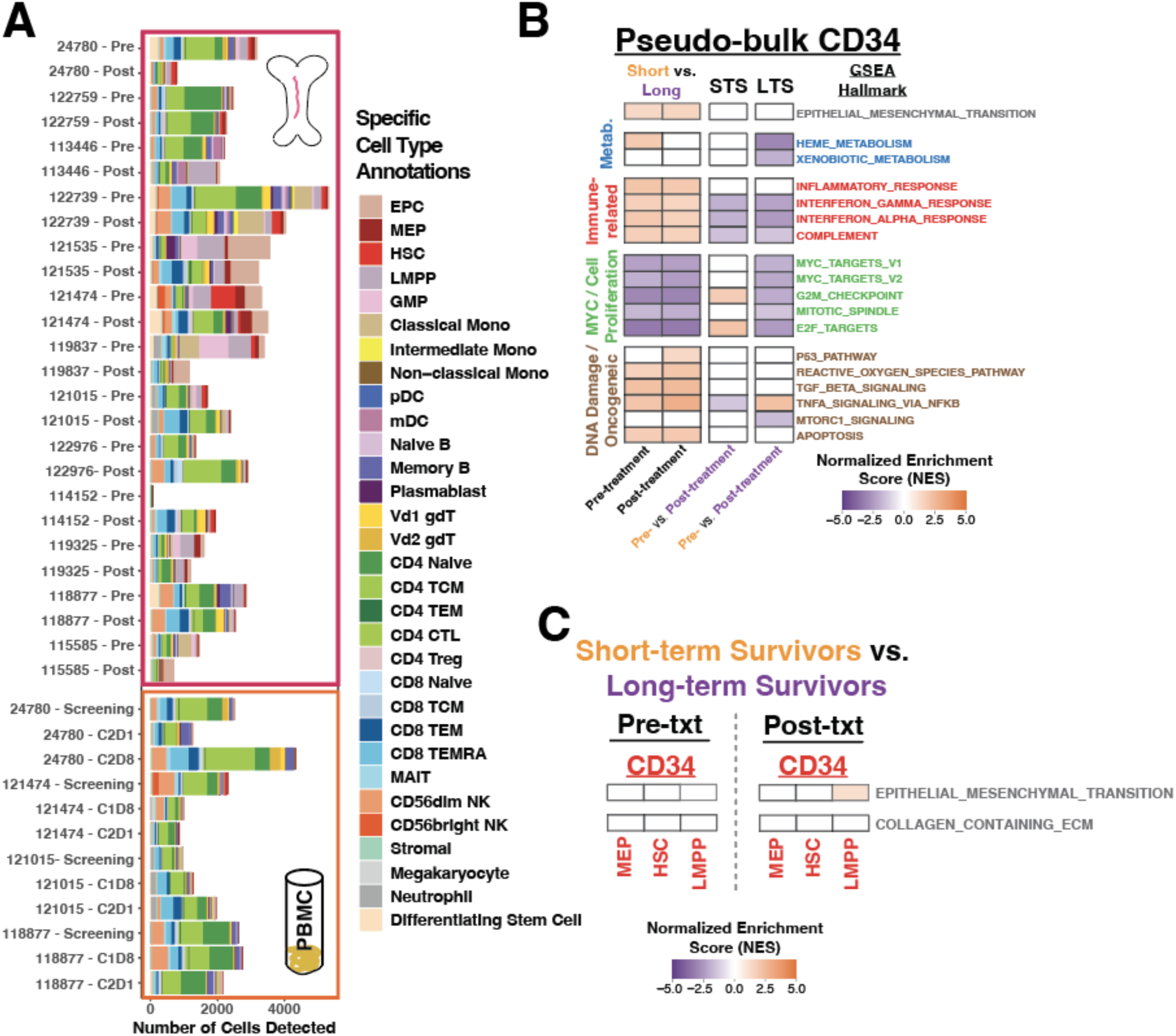
CITE-seq characterizes cellular heterogeneity and transcriptomic dynamics of bone marrow microenvironment cells from MDS patients. **A.** Stacked bar plots illustrating the number of each cell subtypes detected in MDS patient samples across treatment timepoints. **B.** Hallmark GSEA enrichment detected in pseudo-bulk CD34+ bone marrow cells detected by CITE-seq (adjusted p-value < 0.05). **C.** GSEA of EMT and collagen-containing ECM pathways in pseudo-bulk HSPCs in the bone marrow microenvironment (adjusted p-value < 0.05).

**Supplemental Figure 8.**
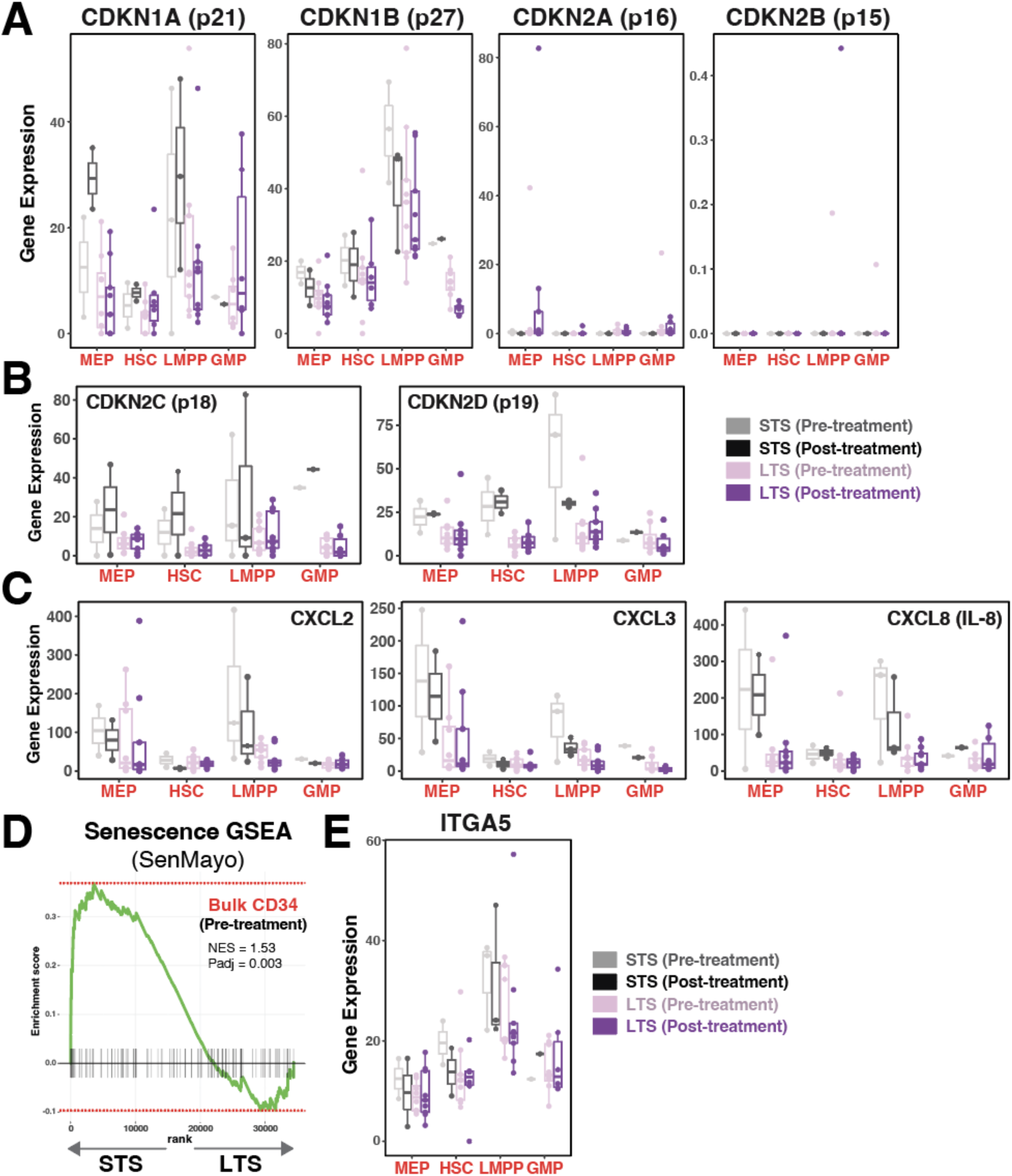
Expression levels of differentially expressed or senescence-associated genes in short survivors’ bone marrow cells. **A-C.** Pseudo-bulk gene expression profiles of non-differentially expressed cyclin dependent kinase inhibitor (CDKN) genes (**A**), or differentially expressed CDKN (**B**) or SASP factors (**C**) in STS- and LTS-HSPCs. **D.** GSEA of senescence-associated pathway, SenMayo, in bulk bone marrow CD34+ cells at pre-treatment timepoint. **E.** Pseudo-bulk expression levels of *ITGA5* detected in HSPCs of STS or LTS bone marrow.

**Supplemental Figure 9.**
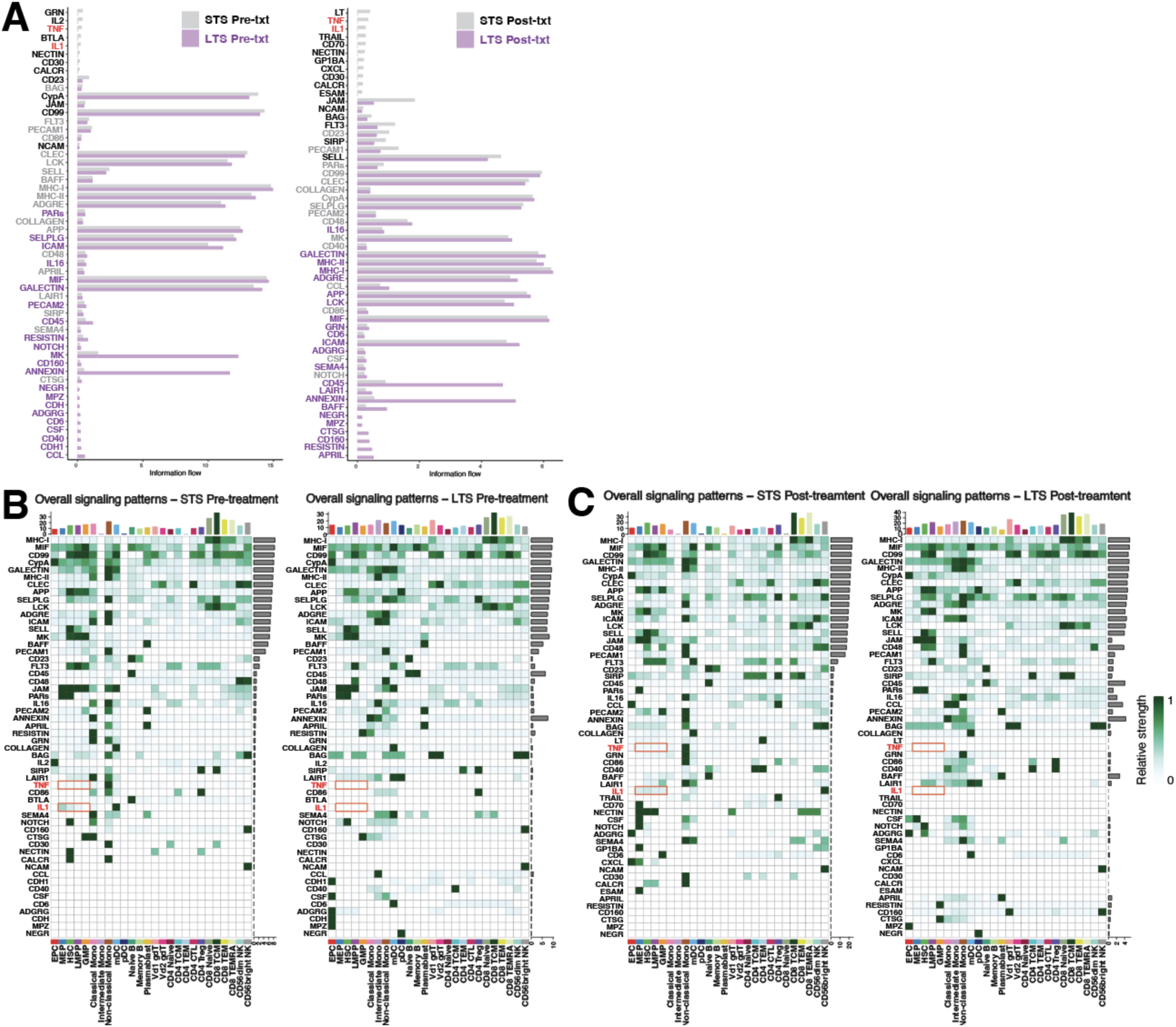
CellChat inference of ligand-receptor interactions across cell subtypes in the bone marrow microenvironment of MDS patients. **A.** Bar plots illustrating preferential information flow for signaling pathways that are detected in STS or LTS bone marrow microenvironment at pre-treatment (left) or post-treatment (right) timepoints. **B-C.** Heatmaps illustrating the relative signaling strength of a signaling pathway contributed by each bone marrow microenvironment cell subtypes from STS (left) or LTS (right) bone marrow at pre-treatment (**C**) or post-treatment (**D**) timepoints.

**Supplemental Figure 10.**
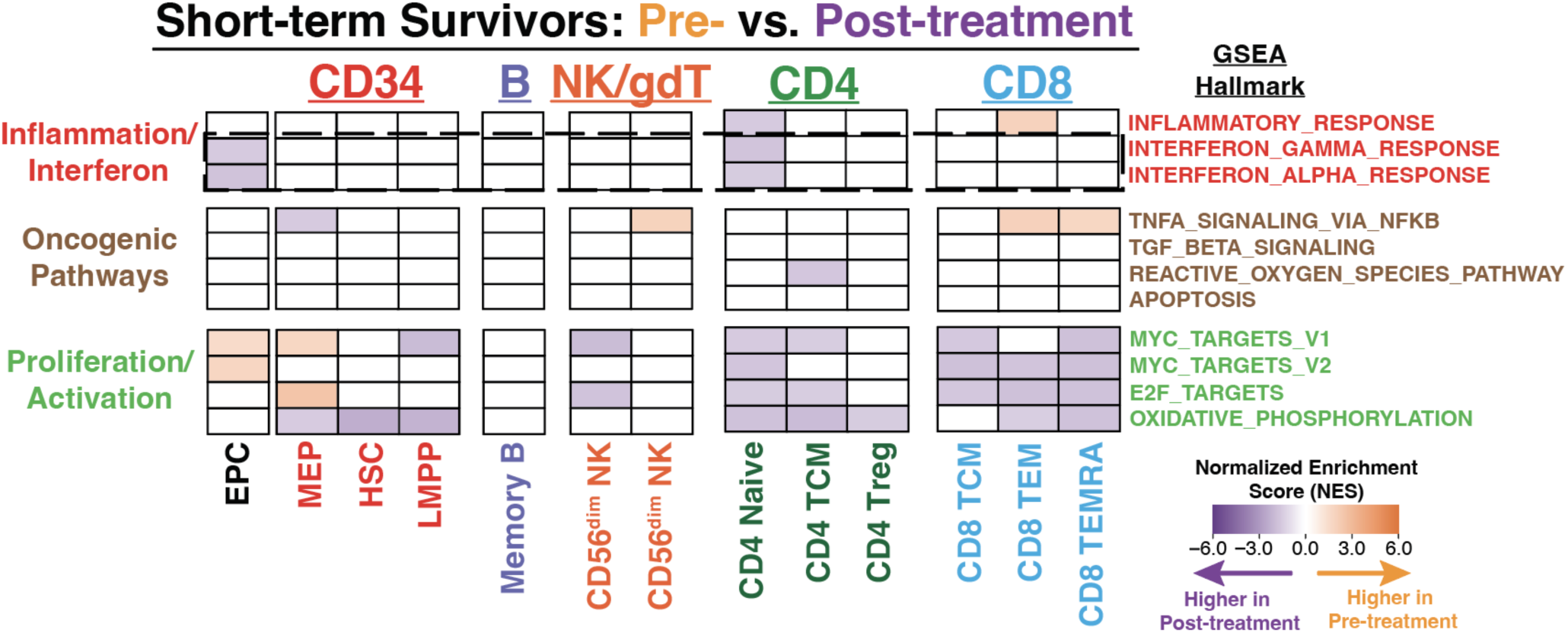
Short survivors’ bone marrow microenvironment lack interferon pathway activation post-treatment. Pseudo-bulk GSEA of hallmark pathways comparing pre-treatment and post-treatment samples of STS cell subtypes (adjusted p-value < 0.05).

**Supplemental Figure 11.**
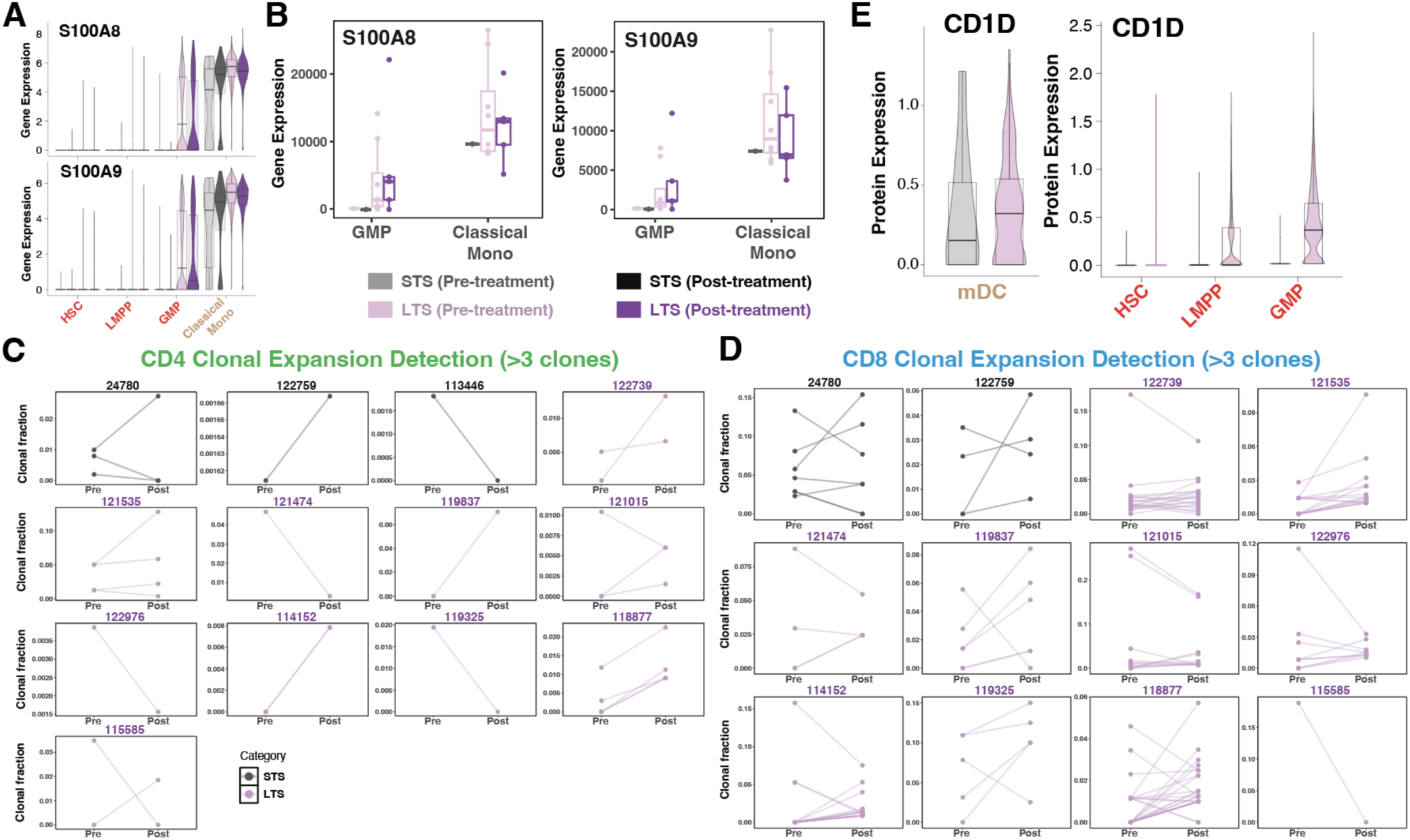
Specific myeloid signatures in long survivors’ bone marrow microenvironment contribute to immune expansion after combination therapy. **A.** Single-cell gene expression distribution of *S100A8/9* in HSPCs and classical monocytes. **B.** Pseudo-bulk expression distribution of *S100A8* and *S100A9* in GMPs and classical monocytes in the bone marrow microenvironment of MDS patients. **C-D.** Clonal expansion detection of CD4 T cells (**C**) and CD8 T cells (**D**) across each MDS patient profiled with CITE-seq. **E.** Single-cell protein abundance distribution of CD1D in mDCs (left) and HSPCs (right).

**Supplemental Figure 12.**
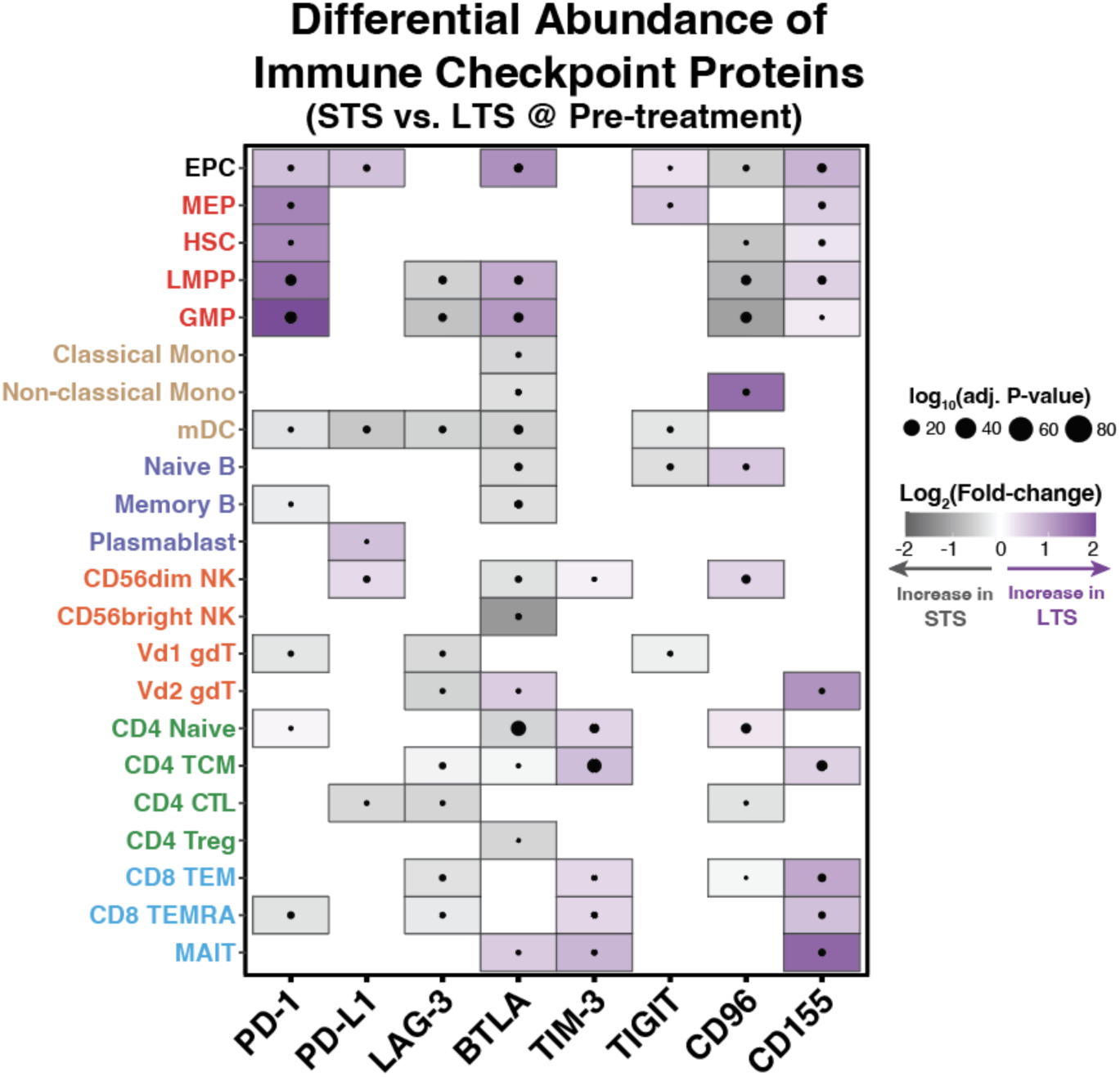
Single-cell differential abundance analysis of immune checkpoint proteins comparing STS or LTS bone marrow microenvironment cell subtypes across pre-treatment timepoint.

**Supplemental Figure 13.**
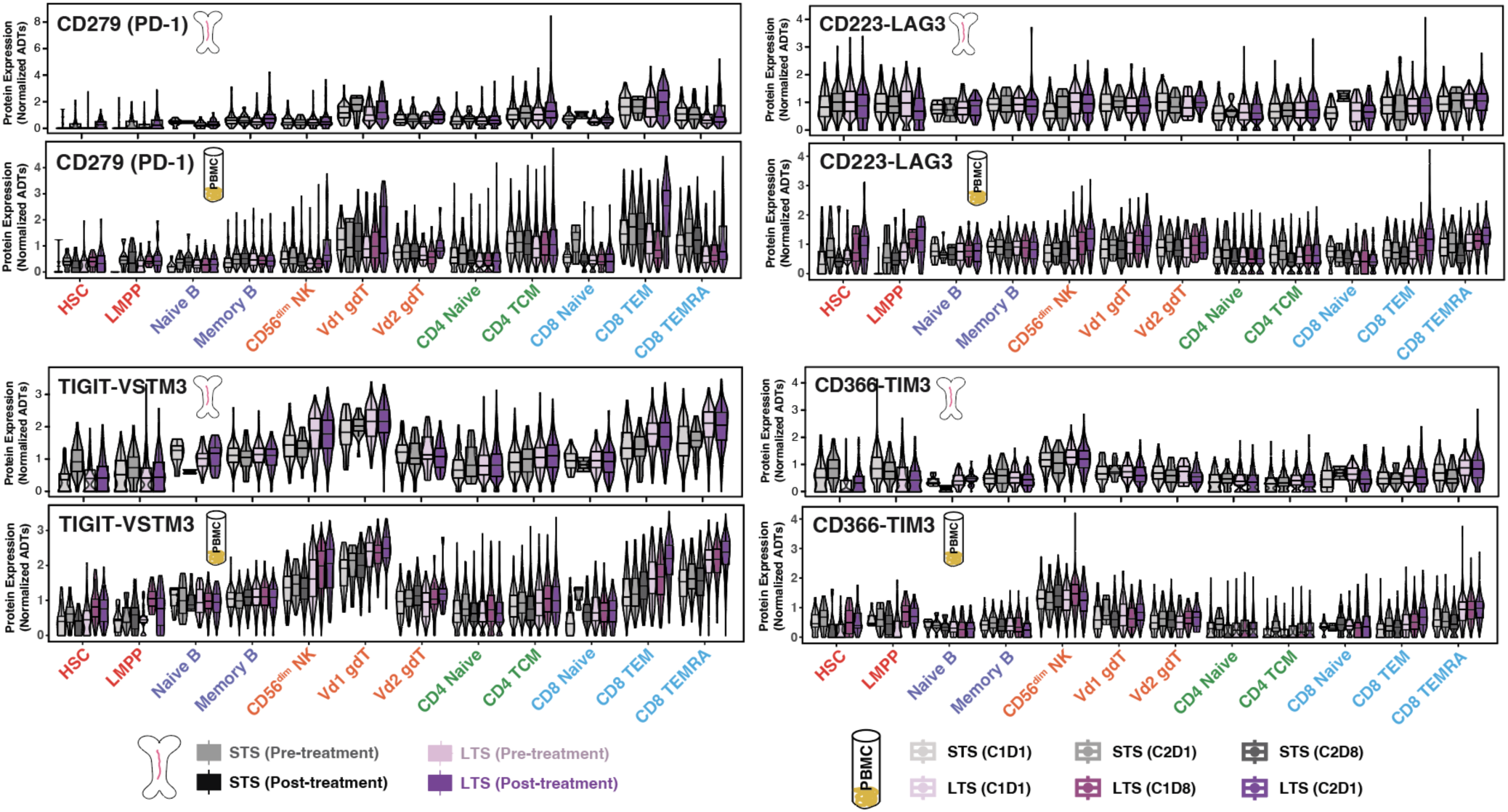
Peripheral blood samples can be valuable surrogates for bone marrow cells to provide prognostic value for combination therapy. Single-cell protein abundance distribution of PD-1, TIGIT, LAG-3, and TIM-3 across cell subtypes detected in both bone marrow (top) and PBMCs (bottom) from MDS patients.

**Supplemental Figure 14.**
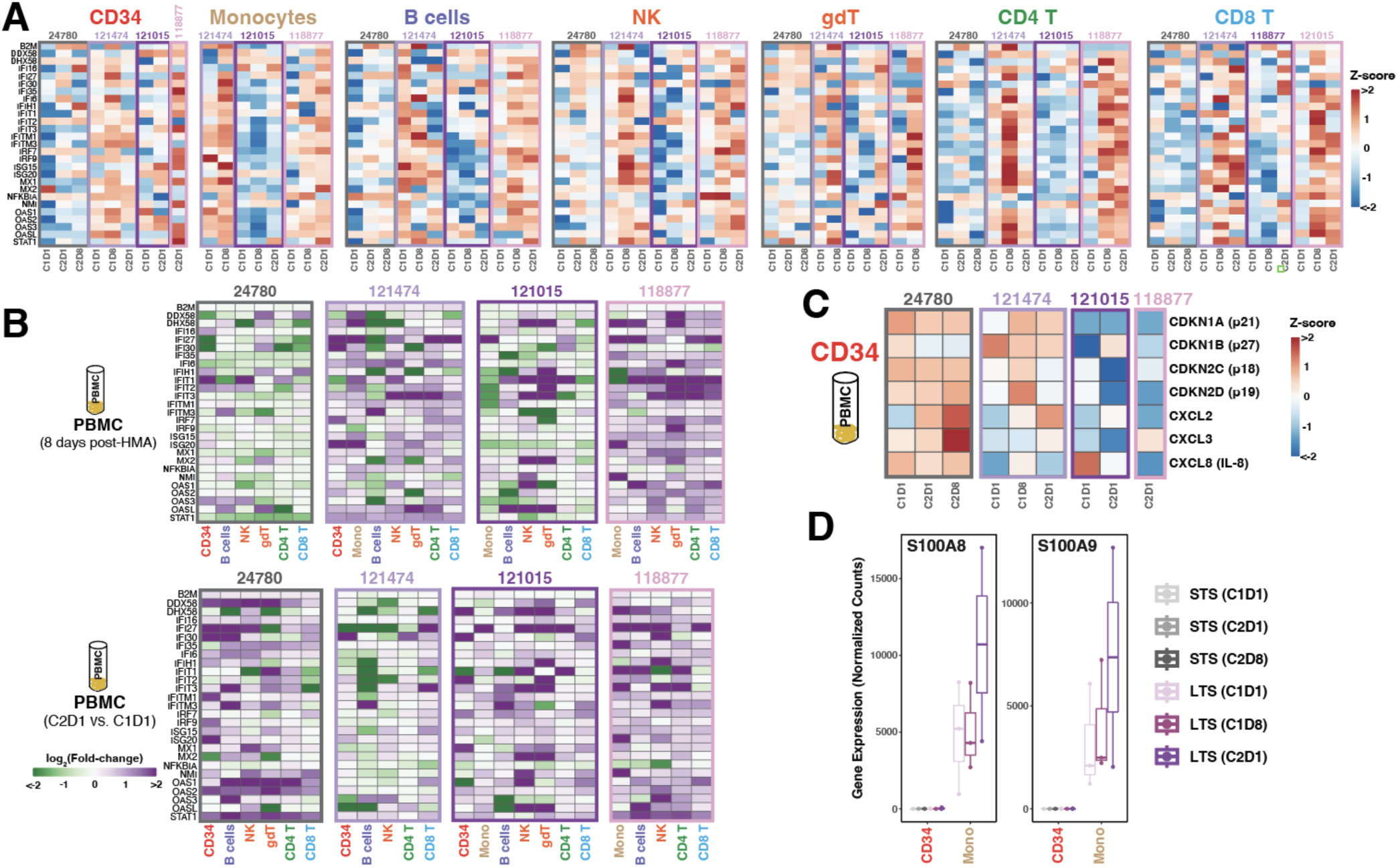
Viral mimicry activation in peripheral blood cells does not associate with patient survival after combination therapy. **A-B.** Heatmaps representing viral mimicry gene expression (**A**) or fold-change (**B**) after combination therapy in pseudo-bulk cell types detected in PBMCs. **C.** Gene expression levels of senescence-associated genes in pseudo-bulk CD34+ cells detected in PBMC samples. **D.** Pseudo-bulk gene expression levels of *S100A8/9* in CD34+ cells or monocytes detected in PBMC samples.

**Supplemental Figure 15.**
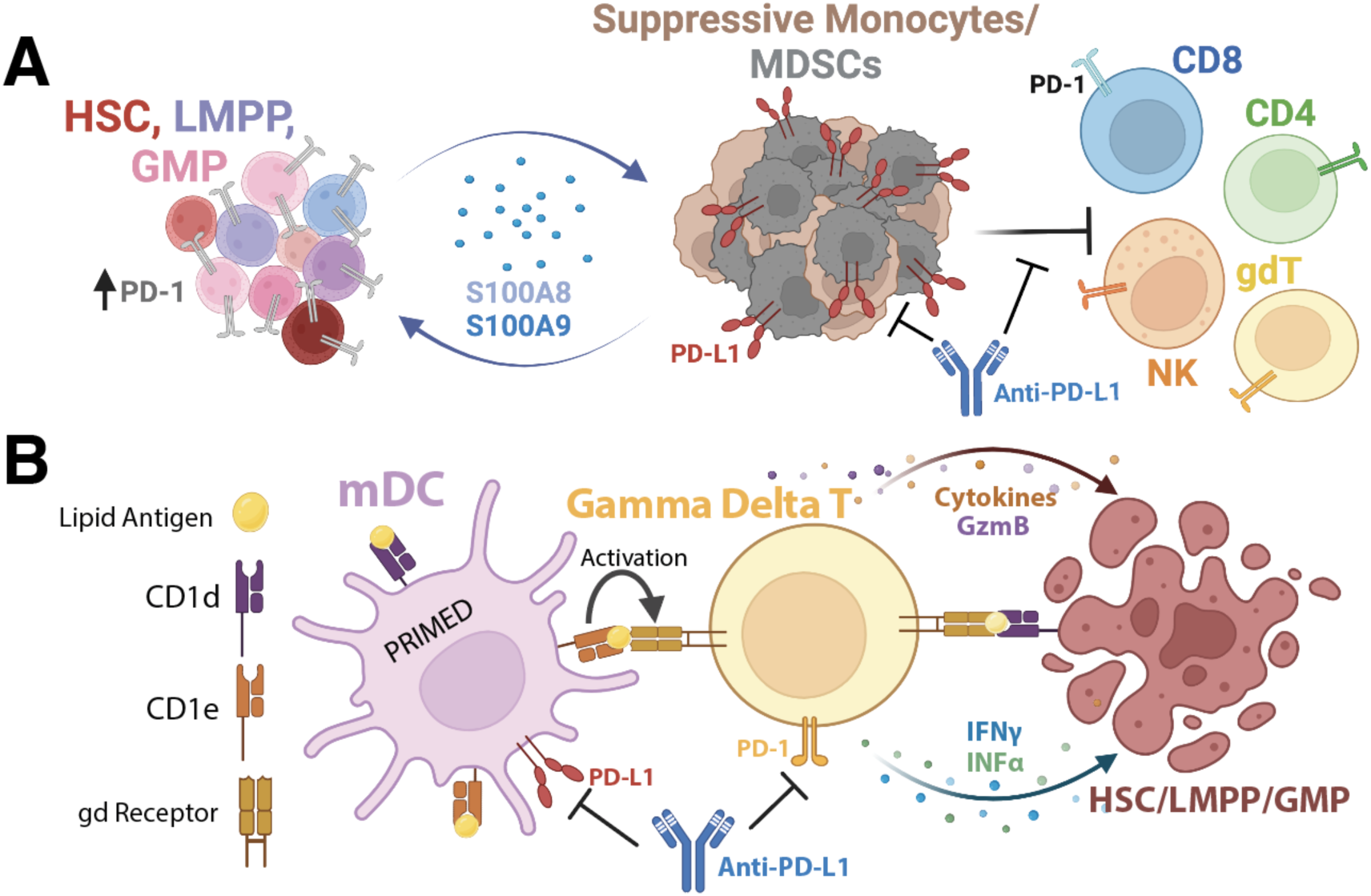
Prognostic immune-mediated mechanisms in LTS bone marrow that contribute to survival after combination therapy. **A.** LTS bone marrow microenvironments show S100A8/9-induced up-regulation of PD-1 on HSPCs and recruitment of suppressive myeloid cells that restrict effector immune function. The combination therapy reduces the presence of suppressive myeloid cells in the LTS bone marrow microenvironment, which associates which global immune activation. **B.** Increased lipid antigen presentation by mDCs in the bone marrow microenvironment can transactivate effector functions in gdT cells to reduce disease burden in long-term survivors. Illustrations were created with BioRender.

## References

1. Volpe, V. O., Garcia-Manero, G. & Komrokji, R. S. Myelodysplastic Syndromes: A New Decade. Clinical Lymphoma, Myeloma and Leukemia vol. 22 Preprint at 10.1016/j.clml.2021.07.031 (2022).

2. Sperling, A. S., Gibson, C. J. & Ebert, B. L. The genetics of myelodysplastic syndrome: From clonal haematopoiesis to secondary leukaemia. Nature Reviews Cancer Preprint at 10.1038/nrc.2016.112 (2017).

3. Corey, S. J. et al. Myelodysplastic syndromes: The complexity of stem-cell diseases. Nature Reviews Cancer Preprint at 10.1038/nrc2047 (2007).

4. Nagata, Y. & Maciejewski, J. P. The functional mechanisms of mutations in myelodysplastic syndrome. Leukemia Preprint at 10.1038/s41375-019-0617-3 (2019).

5. Šimoničová, K. et al. Different mechanisms of drug resistance to hypomethylating agents in the treatment of myelodysplastic syndromes and acute myeloid leukemia. Drug Resistance Updates vol. 61 Preprint at 10.1016/j.drup.2022.100805 (2022).

6. Steensma, D. P. Myelodysplastic syndromes current treatment algorithm 2018. Blood Cancer J 8, (2018).

7. Bewersdorf, J. P. & Zeidan, A. M. Management of patients with higher-risk myelodysplastic syndromes after failure of hypomethylating agents: What is on the horizon? Best Practice and Research: Clinical Haematology vol. 34 Preprint at 10.1016/j.beha.2021.101245 (2021).

8. Jabbour, E. et al. Outcome of patients with myelodysplastic syndrome after failure of decitabine therapy. Cancer 116, (2010).

9. Prébet, T. et al. Outcome of high-risk myelodysplastic syndrome after azacitidine Treatment failure. Journal of Clinical Oncology 29, (2011).

10. Duong, V. H. et al. Poor outcome of patients with myelodysplastic syndrome after azacitidine treatment failure. Clin Lymphoma Myeloma Leuk (2013) doi:10.1016/j.clml.2013.07.007.

11. Roulois, D. et al. DNA-Demethylating Agents Target Colorectal Cancer Cells by Inducing Viral Mimicry by Endogenous Transcripts. Cell 162, 961–973 (2015).

12. Chiappinelli, K. B. et al. Inhibiting DNA Methylation Causes an Interferon Response in Cancer via dsRNA Including Endogenous Retroviruses. Cell 162, 974–986 (2015).

13. Ohtani, H. et al. Activation of a Subset of Evolutionarily Young Transposable Elements and Innate Immunity are Linked to Clinical Responses to 5-Azacytidine. Cancer Res (2020) doi:10.1158/0008-5472.can-19-1696.

14. Ørskov, A. D. et al. Hypomethylation and up-regulation of PD-1 in T cells by azacytidine in MDS/AML patients: A rationale for combined targeting of PD-1 and DNA methylation. Oncotarget (2015) doi:10.18632/oncotarget.3324.

15. Yang, H. et al. Expression of PD-L1, PD-L2, PD-1 and CTLA4 in myelodysplastic syndromes is enhanced by treatment with hypomethylating agents. Leukemia (2014) doi:10.1038/leu.2013.355.

16. Topper, M. J., Vaz, M., Marrone, K. A., Brahmer, J. R. & Baylin, S. B. The emerging role of epigenetic therapeutics in immuno-oncology. Nature Reviews Clinical Oncology Preprint at 10.1038/s41571-019-0266-5 (2020).

17. Jones, P. A., Ohtani, H., Chakravarthy, A. & De Carvalho, D. D. Epigenetic therapy in immune-oncology. Nature Reviews Cancer Preprint at 10.1038/s41568-019-0109-9 (2019).

18. Daver, N. et al. Hypomethylating agents in combination with immune checkpoint inhibitors in acute myeloid leukemia and myelodysplastic syndromes. Leukemia Preprint at 10.1038/s41375-018-0070-8 (2018).

19. O’Connell, C. L. et al. Safety, Outcomes, and T-Cell Characteristics in Patients with Relapsed or Refractory MDS or CMML Treated with Atezolizumab in Combination with Guadecitabine. Clinical Cancer Research 28, (2022).

20. Abbas, H. A. et al. Response to Hypomethylating Agents in Myelodysplastic Syndrome Is Associated With Emergence of Novel TCR Clonotypes. Front Immunol 12, (2021).

21. Stomper, J., Rotondo, J. C., Greve, G. & Lübbert, M. Hypomethylating agents (HMA) for the treatment of acute myeloid leukemia and myelodysplastic syndromes: mechanisms of resistance and novel HMA-based therapies. Leukemia vol. 35 Preprint at 10.1038/s41375-021-01218-0 (2021).

22. Ørskov, A. D. & Grønbæk, K. DNA Methyltransferase Inhibitors in Myeloid Cancer: Clonal Eradication or Clonal Differentiation? Cancer Journal (United States) vol. 23 Preprint at 10.1097/PPO.0000000000000282 (2017).

23. Zhao, G., Wang, Q., Li, S. & Wang, X. Resistance to Hypomethylating Agents in Myelodysplastic Syndrome and Acute Myeloid Leukemia From Clinical Data and Molecular Mechanism. Frontiers in Oncology vol. 11 Preprint at 10.3389/fonc.2021.706030 (2021).

24. Pardoll, D. M. The blockade of immune checkpoints in cancer immunotherapy. Nature Reviews Cancer vol. 12 Preprint at 10.1038/nrc3239 (2012).

25. He, X. & Xu, C. Immune checkpoint signaling and cancer immunotherapy. Cell Research vol. 30 Preprint at 10.1038/s41422-020-0343-4 (2020).

26. Subramanian, A. et al. Gene set enrichment analysis: A knowledge-based approach for interpreting genome-wide expression profiles. Proc Natl Acad Sci U S A 102, (2005).

27. Sidney, L. E., Branch, M. J., Dunphy, S. E., Dua, H. S. & Hopkinson, A. Concise review: Evidence for CD34 as a common marker for diverse progenitors. Stem Cells vol. 32 Preprint at 10.1002/stem.1661 (2014).

28. Novoseletskaya, E. et al. Mesenchymal Stromal Cell-Produced Components of Extracellular Matrix Potentiate Multipotent Stem Cell Response to Differentiation Stimuli. Front Cell Dev Biol 8, (2020).

29. Rusch, R. M. et al. Mscs become collagen-type i producing cells with different phenotype in allogeneic and syngeneic bone marrow transplantation. Int J Mol Sci 22, (2021).

30. Wenk, C. et al. Direct modulation of the bone marrow mesenchymal stromal cell compartment by azacitidine enhances healthy hematopoiesis. Blood Adv 2, (2018).

31. Stoeckius, M. et al. Simultaneous epitope and transcriptome measurement in single cells. Nat Methods (2017) doi:10.1038/nmeth.4380.

32. Stoeckius, M. et al. Cell Hashing with barcoded antibodies enables multiplexing and doublet detection for single cell genomics. Genome Biol (2018) doi:10.1186/s13059-018-1603-1.

33. Hao, Y. et al. Dictionary learning for integrative, multimodal and scalable single-cell analysis. Nat Biotechnol 42, (2024).

34. Hao, Y. et al. Integrated analysis of multimodal single-cell data. Cell 184, (2021).

35. Oetjen, K. A., et al. Human bone marrow assessment by single-cell RNA sequencing, mass cytometry, and flow cytometry. JCI Insight 3, (2018).

36. Hay, S. B., Ferchen, K., Chetal, K., Grimes, H. L. & Salomonis, N. The Human Cell Atlas bone marrow single-cell interactive web portal. Exp Hematol 68, (2018).

37. Love, M. I., Huber, W. & Anders, S. Moderated estimation of fold change and dispersion for RNA-seq data with DESeq2. Genome Biol 15, (2014).

38. Squair, J. W. et al. Confronting false discoveries in single-cell differential expression. Nat Commun 12, (2021).

39. Asada, N. et al. Differential cytokine contributions of perivascular haematopoietic stem cell niches. Nat Cell Biol 19, (2017).

40. Agarwal, P. et al. Mesenchymal Niche-Specific Expression of Cxcl12 Controls Quiescence of Treatment-Resistant Leukemia Stem Cells. Cell Stem Cell 24, (2019).

41. Capone, S. et al. Senescent human hematopoietic progenitors show elevated expression of transposable elements and inflammatory genes. Exp Hematol 62, (2018).

42. Hirai, H., Roussel, M. F., Kato, J.-Y., Ashmun, R. A. & Sherr, C. J. Novel INK4 Proteins, p19 and p18, Are Specific Inhibitors of the Cyclin D-Dependent Kinases CDK4 and CDK6. Mol Cell Biol 15, (1995).

43. Schreiber, M., Muller, W. J., Singh, G. & Graham, F. L. Comparison of the effectiveness of adenovirus vectors expressing cyclin kinase inhibitors p16(INK4A), p18(INK4C), p19(INK4D), p21(WAF1/CIP1) and p27(KIP1) in inducing cell cycle arrest, apoptosis and inhibition of tumorigenicity. Oncogene 18, (1999).

44. Saul, D. et al. A new gene set identifies senescent cells and predicts senescence-associated pathways across tissues. Nat Commun 13, (2022).

45. Fan, G., et al. An immunosuppressive subtype of senescent tumor cells predicted worse immunotherapy response in lung adenocarcinoma. iScience 26, (2023).

46. Jin, S. et al. Inference and analysis of cell-cell communication using CellChat. Nat Commun 12, (2021).

47. Li, X., et al. Inflammation and aging: signaling pathways and intervention therapies. Signal Transduction and Targeted Therapy vol. 8 Preprint at 10.1038/s41392-023-01502-8 (2023).

48. Unnikrishnan, A. et al. Integrative Genomics Identifies the Molecular Basis of Resistance to Azacitidine Therapy in Myelodysplastic Syndromes. Cell Rep 20, (2017).

49. Wang, S. et al. S100A8/A9 in inflammation. Frontiers in Immunology vol. 9 Preprint at 10.3389/fimmu.2018.01298 (2018).

50. Cheng, P. et al. S100A9-induced overexpression of PD-1/PD-L1 contributes to ineffective hematopoiesis in myelodysplastic syndromes. Leukemia 33, (2019).

51. Kapor, S. & Santibanez, J. F. Myeloid-derived suppressor cells and mesenchymal stem/stromal cells in myeloid malignancies. Journal of Clinical Medicine vol. 10 Preprint at 10.3390/jcm10132788 (2021).

52. Kumar, V., Patel, S., Tcyganov, E. & Gabrilovich, D. I. The Nature of Myeloid-Derived Suppressor Cells in the Tumor Microenvironment. Trends in Immunology Preprint at 10.1016/j.it.2016.01.004 (2016).

53. von Wulffen, M. et al. S100A8/A9-alarmin promotes local myeloid-derived suppressor cell activation restricting severe autoimmune arthritis. Cell Rep 42, (2023).

54. Sinha, P. et al. Proinflammatory S100 Proteins Regulate the Accumulation of Myeloid-Derived Suppressor Cells. The Journal of Immunology 181, (2008).

55. Jiang, W., Hu, K., Liu, X., Gao, J. & Zhu, L. Single-cell transcriptome analysis reveals the clinical implications of myeloid-derived suppressor cells in head and neck squamous cell carcinoma. Pathology and Oncology Research 29, (2023).

56. Espinosa-Carrasco, G. et al. Intratumoral immune triads are required for immunotherapy-mediated elimination of solid tumors. Cancer Cell 42, 1202–1216.e8 (2024).

57. Song, L. et al. TRUST4: immune repertoire reconstruction from bulk and single-cell RNA-seq data. Nat Methods 18, (2021).

58. Kadowaki, N. The divergence and interplay between pDC and mDC in humans. Frontiers in Bioscience 14, (2009).

59. McArdel, S. L., Terhorst, C. & Sharpe, A. H. Roles of CD48 in regulating immunity and tolerance. Clinical Immunology vol. 164 Preprint at 10.1016/j.clim.2016.01.008 (2016).

60. Jauch-Speer, S. L. et al. C/EBPδ-induced epigenetic changes control the dynamic gene transcription of S100a8 and S100a9. Elife 11, (2022).

61. Huang, S. et al. CD1 lipidomes reveal lipid-binding motifs and size-based antigen-display mechanisms. Cell 186, (2023).

62. Luoma, A. M., Castro, C. D. & Adams, E. J. γδ T cell surveillance via CD1 molecules. Trends in Immunology vol. 35 Preprint at 10.1016/j.it.2014.09.003 (2014).

63. Kang, T. G. et al. Epigenetic regulators of clonal hematopoiesis control CD8 T cell stemness during immunotherapy. Science (1979) 386, eadl4492 (2024).

64. Duy, C. et al. Chemotherapy induces senescence-like resilient cells capable of initiating aml recurrence. Cancer Discov 11, (2021).

65. Wang, L., Lankhorst, L. & Bernards, R. Exploiting senescence for the treatment of cancer. Nature Reviews Cancer vol. 22 Preprint at 10.1038/s41568-022-00450-9 (2022).

66. Kopylova, E., Noé, L. & Touzet, H. SortMeRNA: Fast and accurate filtering of ribosomal RNAs in metatranscriptomic data. Bioinformatics 28, (2012).

67. Dobin, A. et al. STAR: Ultrafast universal RNA-seq aligner. Bioinformatics (2013) doi:10.1093/bioinformatics/bts635.

68. Tilford, C. A. & Siemers, N. O. Gene set enrichment analysis. Methods Mol Biol 563, (2009).

69. Liao, Y., Smyth, G. K. & Shi, W. FeatureCounts: An efficient general purpose program for assigning sequence reads to genomic features. Bioinformatics 30, 923–930 (2014).

70. Li, H. T. et al. RNA mis-splicing drives viral mimicry response after DNMTi therapy in SETD2-mutant kidney cancer. Cell Rep 42, (2023).

71. Fleming, S. J. et al. Unsupervised removal of systematic background noise from droplet-based single-cell experiments using CellBender. Nat Methods 20, (2023).

72. Amezquita, R. A. et al. Orchestrating single-cell analysis with Bioconductor. Nat Methods 17, (2020).

